# Mitochondrially Transcribed dsRNA Mediates Manganese-induced Neuroinflammation

**DOI:** 10.1101/2025.02.16.638529

**Authors:** Hadassah Mendez-Vazquez, Avanti Gokhale, Maureen M. Sampson, Felix G Rivera Moctezuma, Adriana Harbuzariu, Anson Sing, Stephanie A. Zlatic, Anne M. Roberts, Milankumar Prajapati, Blaine R. Roberts, Thomas B. Bartnikas, Levi B. Wood, Steven A. Sloan, Victor Faundez, Erica Werner

**Affiliations:** Department of Cell Biology, Emory University, 615 Michael St, Atlanta, GA, USA, 30322; Department of Human Genetics, Emory University, 615 Michael St, Atlanta, GA, USA, 30322; Emory Stem Cell and Organoids Core, Emory University, 615 Michael St, Atlanta, GA, USA, 30322; Department of Biochemistry, Emory University, 1510 Clifton Rd, Atlanta, GA, USA, 30322; Department of Neurology, Emory University, 12 Executive Park Dr NE, Atlanta, GA, USA, 30322; Department of Pharmaceutical Sciences, Ferris State University, 220 Ferris Dr, Big Rapids, MI, USA, 49307; George W. Woodruff School of Mechanical Engineering and Parker H. Petit Institute for Bioengineering and Bioscience, Georgia Institute of Technology, 315 Ferst Dr, Atlanta, GA, USA, 30332; Wallace H Coulter Department of Biomedical Engineering, Georgia Institute of Technology, 315 Ferst Dr, Atlanta GA, USA, 30332; Department of Pathology and Laboratory Medicine, Brown University, Providence, RI, USA, 02912

**Author notes:** Corresponding author email: Erica Werner. These authors contributed equally (Hadassah Mendez-Vazquez, Avanti Gokhale).

**Keywords:** dsRNA, manganese, astrocyte, interferon

## Abstract

Manganese is an essential trace element required for various biological functions, but in excess is neurotoxic and leads to significant health concerns. The mechanisms underlying manganese neurotoxicity remain poorly understood. Neuropathological studies of affected brain regions reveal astrogliosis, neuronal loss, and neuroinflammation. Here, we present a novel manganese-dependent mechanism linking mitochondrial dysfunction to neuroinflammation. We found that manganese disruption of the mitochondrial transcriptome processing results in the accumulation of double stranded RNA (dsRNA). This dsRNA is released into the cytoplasm, where it activates the cytosolic sensor MDA5, triggering type I interferon responses and inflammatory cytokine production. This mechanism is evident in 100 day human cerebral organoids, where manganese-increased mitochondrial dsRNA and induced inflammatory responses in mature astrocytes. Similarly, we observed an increase in mitochondrial dsRNA content, the activation of an inflammatory transcriptome and the production of cytokines in female and male mouse brains carrying mutations in the Slc30a10 gene, a model for human hypermanganesemia with dystonia 1 disorder. These findings highlight a previously unrecognized role for mitochondrial dsRNA in manganese-induced neuroinflammation and provide insights into the molecular pathogenesis of manganism. We propose that this mitochondrial dsRNA-induced inflammatory pathway could be active in other neurological diseases caused by environmental or genetic factors.

**Significance Statement:** Environmental exposures and genetic defects that perturb manganese homeostasis are an underappreciated cause of neurodegeneration and neuroinflammation. We describe a new paradigm for inducible neuroinflammation, where manganese disruption of mitochondrial transcriptome processing leads to the accumulation of mitochondrial double-stranded RNA (dsRNA), which activate antiviral responses in the cytoplasm driving type I interferon dependent inflammation. This manganese-dsRNA axis is induced in cell lines in vitro and a subpopulation of mature astrocytes in exposed human cerebral organoids. Brain cortex of mice deficient in the manganese efflux transporter Slc30a10, a genetic model of chronic manganese accumulation, show dsRNA accumulation, and up-regulation of type I interferon response and astrogliosis markers, supporting a role for this pathway in neurotoxicity and parkinsonism.

## Introduction

Manganese is an essential trace metal for living organisms; however, its excess is neurotoxic (Balachandran et al., 2020). Manganese accumulates as a result of environmental exposures, genetic defects, and/or secondary to liver damage (Peres et al., 2016). Contaminated water, air, and soil affect broad human populations while miners, welders, and industrial workers are at occupational exposure risk (Lucas et al., 2015; Kornblith et al., 2018; Schullehner et al., 2020). Mutations in transporters controlling manganese influx (SLC39A14) or efflux (SLC30A10) from cells and organisms increase metal levels in the brain and other organs leading to hypermanganesemia with dystonia 1 and 2 disorders in humans (Mercadante et al., 2019; Taylor et al., 2019; Rodichkin and Guilarte, 2022).

In the brain, manganese preferentially accumulates in the basal ganglia, causing manganism, which manifests with neuropsychiatric symptoms, and then progresses to extrapyramidal dysfunction and dystonia (Lucchini et al., 1999; Anagianni and Tuschl, 2019). Chronic low-level exposure increases the risk of cognitive, neuropsychiatric and behavioral symptoms in young and at-risk adult individuals (Mergler et al., 1999; Schullehner et al., 2020).

The mechanisms for manganese brain toxicity remain poorly understood. Neuropathology of the globus pallidus in non-symptomatic and symptomatic individuals following manganese exposures, showed astrogliosis, increased glia counts and mild neuronal loss (Olanow, 2004; Perl and Olanow, 2007; Vezer et al., 2007; Gonzalez-Cuyar et al., 2014). Microglia and astrocytes release cytokines at early and late stages of disease in manganese exposed mice, indicating a neuroinflammatory component to the pathogenic mechanism of toxicity in the brain (Antonini et al., 2009; Filipov and Dodd, 2012; Tjalkens et al., 2017; Fan et al., 2020; Soto-Verdugo and Ortega, 2021). Both glial cell types release cytokines in vitro (Sidoryk-Wegrzynowicz and Aschner, 2013), with microglia derived factors enhancing astrocyte responses (Kirkley et al., 2017). These glial interactions exacerbate neuronal toxicity (Zhang et al., 2009; Popichak et al., 2018). Neuroinflammation is a well-established response to injury and is proposed to be disease-specific and involved in both, disease initiation and progression (Frank-Cannon et al., 2009; Glass et al., 2010). Here, we present a novel mechanism for manganese induced neuroinflammation triggered by the generation of endogenous double stranded RNA (dsRNA) in mitochondria.

We previously showed that manganese disrupts processing of the mitochondrial transcriptome and induces accumulation of un-processed mitochondrial transcripts intermediaries (Werner et al., 2022). Processing of the mitochondrial polycistronic RNA is necessary to generate functional ribosomal, transfer, and messenger RNAs, which are required to synthetize the thirteen mitochondrial-encoded proteins necessary for the assembly and function of the respiratory chain (D’Souza and Minczuk, 2018). The transcription of both strands of mitochondrial DNA generates long complementary RNA which are homeostatically regulated by mitochondrial specific post-transcriptional modifications and degradation of non-coding transcripts (Murphy et al., 1975; Santonoceto et al., 2024). Accumulation of complementary RNA has the potential to generate mitochondrial dsRNA, which can translocate to the cytoplasm, activate cytosolic sensor molecules and trigger antiviral signaling pathways to produce type I interferon (IFN) responses and inflammation (Barshad et al., 2018; Chen and Hur, 2022; Grochowska et al., 2022). We hypothesized that manganese-induced transcriptome defects are sufficient to induce inflammatory signaling. We demonstrate that sub-cytotoxic manganese levels induce mitochondrial dsRNA, which triggers the IFN type I response and cytokine production, in cell lines and in the most mature subpopulation of astrocytes in 100-day old human cerebral organoids. These phenotypes are reproduced in brains of the Slc30a10 mutant mice, a genetic model of manganese accumulation (Quadri et al., 2012; Tuschl et al., 2012).

## Materials and Methods

### Experimental Design and Statistical Analyses

We employed two cell lines and used genetic and pharmacological tools to dissect effectors in the mitochondrial dsRNA-induced signal transduction pathway. To demonstrate manganese induction of these pathways in disease relevant tissues, we carried out studies in human cerebral organoids, hiPSC derived astrocytes, and tissues of the Slc30a10^-/-^ mouse. Critical experimental variables for replication, sample size and statistical test employed are reported in the figure legends.

### Mice

Mice were group housed in a standard 12 h light/dark cycle and were fed Rodent Diet 20 (cat# 5053, PicoLab) *ad libitum*. All animal care protocols and procedures were followed in accordance with NIH and with approval of Emory University’s Institutional Animal Care and Use Committee. Female and male Slc30a10^-/-^ mice in C57BL/6N background were genotyped as in (Mercadante et al., 2019) and punch biopsies from cortex and liver were obtained as before (Zlatic et al., 2023). The samples used in experiments shown in Figure 1 and 6, were generously provided by Dr. Bartnikas.

**Figure 1:**
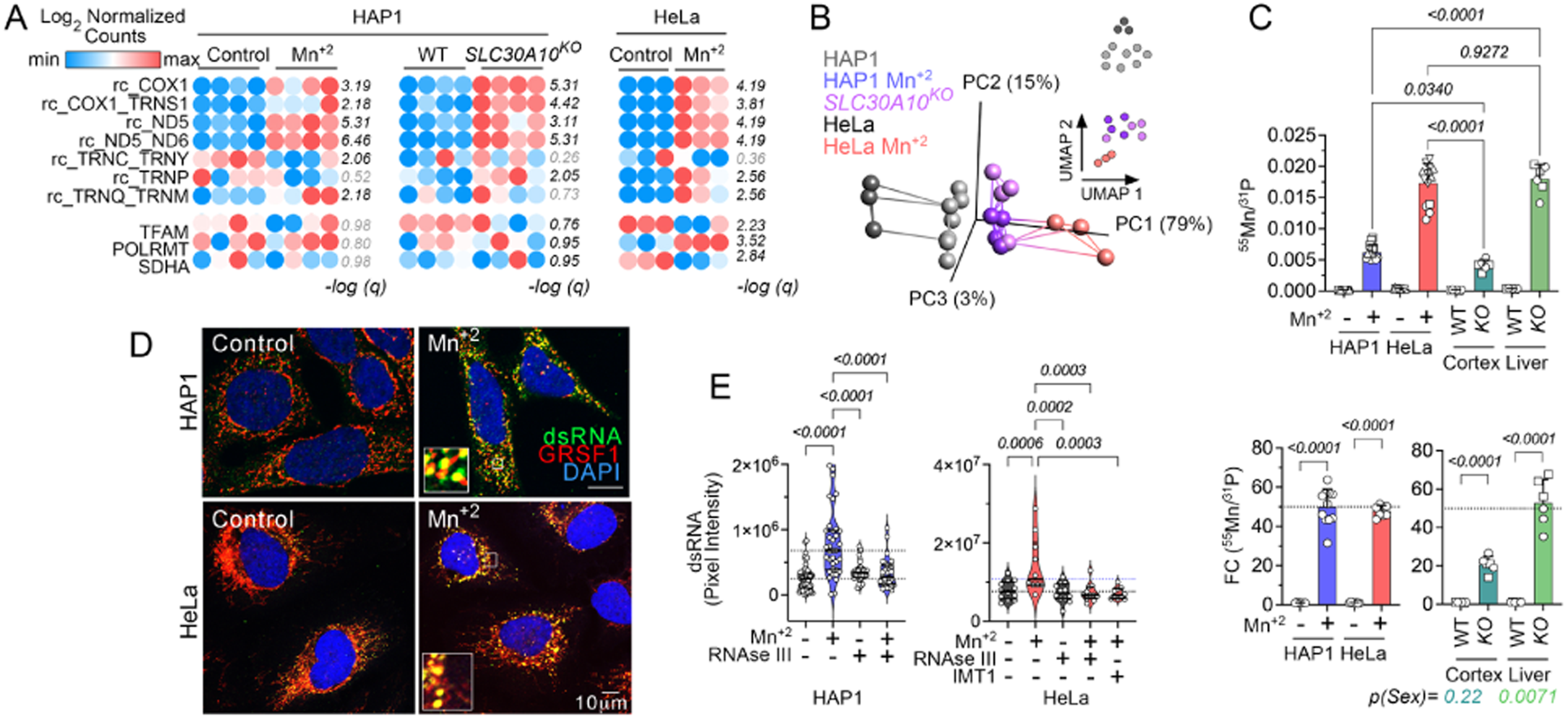
Manganese induces the accumulation of noncoding mitochondrial RNA sequences and mitochondrial dsRNA. a) Heatmaps of log_2_ CLTC normalized MitoString counts of RNA hybridized with probes targeted to non-coding sequences in the light strand transcripts and for nuclear encoded mitochondrial proteins as control. RNA was isolated from four replicate samples of either wild-type HAP1 or HeLa treated either in the absence or presence of manganese (300μM and 800μM, respectively) for 24h, or untreated wild-type and SLC30A10 null HAP1 cells. Numbers in italics represent −log10 p values (t-test followed by Benjamini-Hochberg FDR correction, *q*). b) Manganese treatment and SLC30A10 deficiency similarly alter the MitoString transcriptome. Linear and non-linear data reduction by principal component analysis (PCA) and Uniform Manifold Approximation and Projection (UMAP), respectively of all the RNA counts measured by the MitoString panel. Lines connecting PCA represent Euclidean distances. c) Top Panel: Manganese enrichment reflected by metal levels normalized by phosphate content measured by ICP-MS in HAP1 or HeLa cell pellets at 24h post treatment in at least 3 independent experiments represented by different symbols, and in brain cortex punch biopsies or liver of 8 weeks old wild type (WT) and Slc30a10^-/-^ ( 3 female (○), 3 male (□). No sex differences were noted in brain following one-way ANOVA (treatment: F (7, 76) = 249.8) followed by Šídák’s multiple comparisons test across samples. Bottom Panel: Fold of change of the metal content normalized by phosphate shown in (a), where data from two experiments are combined and in wild type (WT) and Slc30a10^-/-^ mouse cortex and liver. Two-way ANOVA (cortex: genotype F (1, 8) = 149.1; sex: F (1, 8) = 1.748) Liver: genotype: F (1, 8) = 358.7; sex: F (1, 8) = 12.84) followed by Šídák’s multiple comparisons test. d) Immunofluorescence detection of dsRNA and the mitochondrial RNA granule component, GRSF1, in HAP1 and HeLa cells treated either in the absence or presence of manganese. e) Quantification of dsRNA overlapping with GRSF1 in cells treated either in the absence or presence manganese, the POLRMT inhibitor IMT1 (1µM), or with dsRNA specific RNAse III before addition of the dsRNA J2 antibody. Dotted lines represent the average intensity of cells treated in the absence or presence manganese. One-way ANOVA (HAP1 F (5, 216) = 24.76: HeLa: F (4, 50) = 7.243) followed by Šídák’s multiple comparisons tests.

### Cell lines and media

HAP1 wild type and SLC30A10^-/-^ cells were grown in Iscove’s DMEM 10%FBS at 37°C and 5%CO_2_. HeLa cells were grown in DMEM 10%FBS 10mM Glutamine at 37°C and 5%CO_2_. Human iPSC TMOi001-A (Gibco cat #A18945) were obtained through Emory Stem Cell and Organoids Core. This iPSC line was developed by reprogramming CD34+ human umbilical cord blood cells using an episomal system to deliver seven-factors: SOX2, OCT4 (POU5F1), Kruppel-like factor 4 (KLF4), MYC (myelocytomatosis oncogene), NANOG (homeobox transcription factor), LIN28, and simian vacuolating virus 40 large (SV40L) T antigen. iPSCs were cultured in feeder-free conditions in dishes coated with Embryonic Stem Cell qualified Matrigel (Corning). iPSCs were negative for mycoplasma, passaged using ReLeSR (Stem Cell Technologies), cultured in StemFLex media (Thermo Fisher Scientific) and incubated at 37°C and 5% CO_2_. Astrocytes were differentiated and matured from neuroprogenitor cells employing StemCell differentiation and maturation media.

Cerebral organoids were developed as previously described (Lancaster et al., 2017). On the first day, iPSCs were dissociated using Accutase (Stem Cell Technologies) and 18,000 cells were seeded for each organoid on 5-10 fiber scaffolds in low attachment plates. After 5 days, embryoid bodies were formed and transferred to induction media (Stem Cell Technologies) to direct them towards neuronal fate. After another 2 days, cells were expanded in Matrigel droplets for 3 days, then moved to organoid maturation media (Stem Cell Technologies). After 2 weeks in culture, the Matrigel was removed from the organoids by mechanical dissociation and the organoids were grown on an orbital shaker. Maturation media was changed twice a week during organoids growth for up to 100 days in culture.

### Transfections and luciferase measurements

Stable cell lines were generated by transfection of HAP1 and HeLa cells with Lipofectamine 3000 (Invitrogen) and plasmid pNiFty3-L-Fluc-Puro (InvivoGen) followed by selection in media containing 5μg/ml Puromycin (Gibco). High Molecular Weight Poly I:C (InvivoGen) was delivered at 1μg/ml by direct addition to the media at the time of manganese treatment. The Sting agonist 2’3’ cdAM(PS)2 (ADU-S100) was delivered by transfection with Lipofectamine 3000. Following 24h treatment, the cells were washed with PBS, incubated with luciferase substrate Bright Glo (Promega) for 10minutes at room temperature and luminescence measured in a white plate. SiRNA (Dharmacon) was delivered at 50nM by two sequential reverse transfections (at day 1 and 3) with RNAi-Max (Invitrogen). Cells were treated on day 5 post-transfection and analyzed at day 6. Percentage of inhibition was calculated using as a reference the fold of change induced by manganese treatment in scrambled siRNA transfected cells.

### Immunoblot

Cells were washed with cold phosphate-buffered saline and lysed in 10 mM HEPES pH 7.4, 150 mM NaCl, 1 mM ethylene glycol-bis(β-aminoethylether)-*N*,*N*,*N*′,*N*′-tetraacetic acid (EGTA), and 0.1 mM MgCl2, with 0.5% Triton X-100 and Complete anti-protease. Proteins were separated by SDS-PAGE in 4-20% gels and transferred using the semidry transfer method to polyvinylidene difluoride membranes. Membranes were blocked in 5% nonfat milk and incubated overnight with primary antibody in 3% bovine serum albumin at 4°C. Detection HRP labelled secondary was diluted 1:5000 and activity was detected with Western Lightning Plus ECL substrate (PerkinElmer, NEL105001EA) and exposure to GE Healthcare Hyperfilm ECL.

### Immunofluorescence

Cells were fixed in 4% PFA at 37°C for 20 minutes and permeabilized with 0.25% triton X100 on ice for 5 minutes. For dsRNA detection, permeabilization included 20U/ml SUPERase-In RNAse inhibitor. Control wells were treated with 5U/ml RNAseIII in PBS with 10mM MgCl_2_ for 30 min at 37°C. After blocking, primary antibodies were added: J2 1:300, GRSF1 1:500, Tom20 1:300, GFAP 1:300. Slides were imaged in a Nikon A1R HD25 confocal microscope with a 60x Apo lens (NA=1.4,WD=140uM). dsRNA localized to the mitochondrial granule positive for GRSF1 was quantified with Image J/Fiji software in the focal plane of maximal intensity of a Z-stack. On both images, background was subtracted with a 30 radius and segmented by thresholding. Measurements of fluorescence of dsRNA per cell were directed and limited to the corresponding GRSF1 image. At least 30 cells in a minimum of 5 fields were measured. Organoids were fixed overnight in 4%PFA and then incubated in sucrose before embedding. 8μm paraffin sections were re-hydrated by alcohol dilution series washes and antigen retrieval was performed for 20 min in Citrate buffer pH6 with 0.05% Tween at 60% power in the microwave and then stained as described above. Organoid images were corrected by brightness and contrast to distinguish positive from negative cells and background signal. Cells within 300μm from the organoid surface, in 5 different captured fields were identified as GFAP positive and then classified as dsRNA positive or negative. Identical scoring was obtained by two independent observers. Eight-week old Slc30a10^-/-^ mice and wild type littermates were anesthetized with 1% pentobarbital and transcardially perfused with ice cold PBS containing heparin followed by ice cold 4% paraformaldehyde. Brains were removed and post-fixed with 4% paraformaldehyde for 24h and subsequently dehydrated in 30% sucrose in PBS for 48h. Tissues were embedded in optimal cutting temperature compound (OCT Compound, SAKURA, USA) and sectioned coronally (12 μm) by cryostat microtome (CM3050S, Leica, Germany). For immunofluorescence staining, the sections were blocked with 5% normal goat serum with 0.3% Triton X-100 and Fc receptor blocker (Miltenyi) and 20U/ml SUPERase-In RNAse inhibitor for 1h at room temperature and then incubated with primary antibodies for 2h at room temperature. Control sections were incubated with 5U/ml RNAseIII in PBS with 5mM MgCl2 for 1h at 37°C before primary antibody. Endogenous peroxidases were inhibited with 3% H_2_O_2_ for 30 minutes. The J2-dsRNA signal was increased by tyramide amplification (Vector Lab). Following anti mouse HRP incubation for 1h, tyramide was applied for 5min and followed by incubation with the appropriate fluorescent secondary antibodies for 2h at room temperature. Ten fields per section were captured as z-stacks and processed using ImageJ/Fiji software. Amplified J2 fluorescence intensity was quantified in astrocytes by overlying a mask of GFAP staining generated from maximal intensity projection of 3 consecutive z-slices to increase the quantified area and in neurons by overlying the TOM20 signal of Nissl positive cells in a single z-slice.

### Cytokine measurements

Human Cytokine Array were used to profile cytokines in conditioned media following 48h manganese treatment in HeLa (750μM) and HAP1(150μM) cells. A high and a low exposure film was quantified by thresholding and measuring the signal intensity with the circular brush selection in FIJI/Image J on each spot and an equal adjacent area to subtract local background. CXCL8/IL8 was measured in media conditioned for 48h employing Elisa assays following manufacturer’s protocols. The human Proinflammatory 9-Plex panel from MesoScale and R-PLEX Human B2M Antibody Set were employed to measure cytokines and beta 2 microglobulin in conditioned media from organoid treated for 48h by the Emory Multiplexed Immunoassay Core.

### Cell survival assay

Cells were plated at 2,000 cells/well the day before drugs were added. Cells were treated with the drug concentrations indicated for 72h and 10% Alamar blue in media (Resazurin) was incubated for 2 hours at 37°C.Fluorescence excitation at 530–570 nm and emission maximum at 580–590 nm was measured in a microplate reader (BioTek Synergy HT). Percent survival was calculated by subtracting the background of Alamar blue and normalizing to the untreated condition for each genotype. Individual data points represent the average survival of triplicate treatments for each concentration.

### Total RNA extraction and NanoString mRNA Quantification

Cells were solubilized in 1 ml of TRIzol (Invitrogen). RNA Extraction and NanoString processing was done by the Emory Integrated Genomics Core. RNA quality was assessed by bioanalyzer. mRNA counts were normalized to housekeeping genes TBP for the neuroinflammatory panel and CLCT for the MitoString using nSolver software. Normalized data were further processed and visualized in Qlucore.

### Cytokine /Chemokine Multiplex Luminex Immunoassay

Frozen mouse brain samples were provided by Dr. Tom Bartnikas. Shavings of frozen tissue were collected and lysed in 8M urea in 100mM PO_4_ with Complete anti-protease and PhosSTOP phosphatase inhibitor and homogenized by sonication (Fisher Scientific, Sonic Dismembrator Model 100). Protein concentration was measured in triplicate using the Pierce BCA Protein Assay Kit (Thermo Fisher Scientific) according to manufacturer protocol. 5 µg of brain and 15 µg of liver tissue samples were diluted using urea buffer and analyzed using the Milliplex MAP Mouse Cytokine/Chemokine Multiplex assay (Millipore Sigma, St. Louis, MO, USA, MCYTMAG-70K-PX32) Eotaxin/CCL11, G-CSF, GM-CSF, IFN-γ, IL-1α, IL-1β, IL-2, IL-3, IL-4, IL-5, IL-6, IL-7, IL-9, IL-10, IL-12 (p40), IL-12 (p70), IL-13, IL-15, IL-17, IP-10, KC, LIF, LIX, MCP-1, M-CSF, MIG, MIP-1α, MIP-1β, MIP-2, RANTES, TNF-α, VEGF. Assays were read out with a MAGPIX Luminex instrument (Luminex, Austin, TX, USA).

### Metal measurement by ICP mass spectrometry

Procedures to measure manganese, iron, zinc, calcium and phosphate were described previously (Lane et al., 2022). Briefly, cells were detached with trypsin and pelleted at 800 × g for 5 min at 4°C and digested by adding 50 µL of 70% trace metal basis grade nitric acid with heating at 95°C for 10 min. Metal levels were quantified with a triple quad ICP-MS instrument (Thermo Fisher, iCAP-TQ) operating in oxygen mode under standard conditions (RF power 1550 W, sample depth 5.0 mm, nebulizer flow 1.12L/min, spray chamber 3C, extraction lens 1,2 −195, –-15 V). Oxygen was used as reaction gas at 0.3 mL/min to remove polyatomic interferences and mass shift target elements. External calibration curves ranging from 0.5 to 1000 µg/L were generated for each element with a multielemental standard (ICP-MSCAL2-1, AccuStandard, USA). Scandium (10 µg/L) was used as internal standard diluted into the sample in-line. Samples were introduced into the ICP-MS using the 2DX PrepFAST M5 autosampler (Elemental Scientific) equipped with a 250 µL loop and using the 0.25 mL precision method following manufacturer recommendations. Serumnorm (Sero, Norway) was used as a standard reference material, yielding values for elements of interest within 20% of the accepted value. Quantitative data analysis was conducted with Qtegra software and exported to Excel for further statistical analysis.

### Cell isolation and capture for single cell RNAseq

Organoids were dissociated into a single cell suspension using previously published methods (Zhang et al., 2016) adapted for organoids dissociation by decreasing volumes and omission of filtering steps. Briefly, organoid tissue was chopped using a scalpel and digested in 10ml with 25 U/ml papain (Worthington) and 12.5 U/ml DNAseI (Worthington) in a 5% CO2 incubator at 34°C on an orbital shaker at 123rpm for 45 min. Cells were washed with a protease inhibitor solution (ovomucoid). Following addition of Ovomucoid, the tissue was further triturated by pipetting and once single cell suspensions achieved, the cells were collected by centrifugation. All dissociation and trituration steps were performed in the presence of Protector RNAse inhibitor, transcription inhibitor actinomycin D, and translation inhibitor anisomycin based on published recommendations (Marsh et al., 2022). Up to 10,000 cells per organoid were captured with a Chromium GEM-X-Single Cell 3’ Chip kit v4 from 10X Genomics. Libraries were prepared using Chromium Next GEM Single Cell 3’ Kit v4 with dual index labeling and sequenced on a standard Illumina platform.

### Single-cell sequencing analysis

The sequencing data were aligned to the human genome and demultiplexed using Cell Ranger. Analysis was performed in R Studio using Seurat. For quality control, cells with <8% mitochondrial genes were excluded, and counts were filtered for 500-8000 range. Cells were integrated and normalized using SCT transform to generate a UMAP and clustering based on based on shared nearest neighbors. All featureplots, dot plots, and differential expression are based on non-transformed RNA counts. Differential expression was performed using pseudobulk data aggregation and Findmarkers DeSEQ2. Heatmaps are based on scaled RNA counts data.

### Statistical analyses

NanoString data was normalized with nCounter and then processed with Qlucore Omics Explorer Version 3.6 Software. ANOVA and paired analyses were conducted with Prism Version 10.2.2. Gene ontology studies were performed with ENRICHR (Xie et al., 2021).

Luminex Cytokine Statistical and Multivariate Analyses, Wilcoxon rank sum test was performed in R. Partial least squares regressions (PLSRs) and discriminant PLSRs (D-PLSRs) were performed in R using the *ropls* package v1.4.2. The data were z-scored before being input into the function. Cytokine measurements were used as the independent variables, and the discrete regression variable in all D-PLSR analyses was genotype/sex. Error bars for LV loadings were calculated by iteratively excluding K samples without replacement 100 times (leave-K-out cross-validation, LKOCV) and regenerating the D-PLSR model each time. Error bars in the LV1 plots report the mean and SD computed across the models generated to provide an indication of the variability within each cytokine among the models generated.

Statistical analysis of dsRNA quantification in brain was performed with the SuperPlots tool (Lord et al., 2020)

## Results

### Exposure to manganese leads to the accumulation of mitochondrial dsRNA

We previously found that manganese alters mitochondrial RNA processing in HAP1 and SH-SY5Y, measured by molecular counting employing a custom NanoString panel designed for multiplexed detection of mitochondrial mRNA, tRNA-mRNA, and tRNA-rRNA junctions as well as non-coding regions in the polycistronic RNA arising from the transcription of both, the heavy (H) and light (L) strand of mtDNA (Wolf and Mootha, 2014; Werner et al., 2022). We observed the accumulation of non-coding RNA sequences of the L strand transcript, which are normally rapidly degraded (Barshad et al., 2018; Santonoceto et al., 2024), in HAP1 SLC30A10^ko^cells as well as HAP1 and HeLa cells treated for 24h with manganese at 300 and 800µM respectively (Fig. 1A). In contrast, transcripts levels for TFAM, SDHA, or POLRMT did not change consistently, indicating no change in mitochondrial mass. Linear and non-linear dimensionality reduction of the data by principal component analysis and by Uniform Manifold Approximation and Projection (UMAP, Fig. 1B), demonstrated clustering of all the control samples and segregation away from all samples with increased manganese resulting from either manganese exposure or mutagenesis of the SLC30A10 efflux transporter (Fig. 1B). These were the most robust and consistent changes among both cell types (Fig. S1A). To assess possible mitochondrial RNA processing mechanisms disrupted by manganese exposure, we compared the manganese transcriptome responses with those reported by Wolf et al. after screening of mitochondrial RNA binding proteins (Wolf and Mootha, 2014). The manganese exposure mitochondrial transcriptome did not resemble any of the Wolf’s reported patterns (Fig. S1A). These include the cleavage activity either at 3’ or 5’ tRNA-gene junctions, which can be estimated by changes in the mRNA/junction ratios (Fig. S1B), the downregulation of SUPV3L, which Wolf reported causes accumulation of RNA from the L strand, with concurrent accumulation of ND1, ND2 and ND3 transcripts or the activity of PNPT1 or MTPAP (Fig. S1A).

The alterations of the mitochondrial transcriptome after manganese exposure of wild-type cells occur at IC_50_ metal doses (50% death when survival was measured at day 3 post-manganese treatment ((Werner et al., 2022), Fig. 3E and Fig. S2C). These treatment doses resulted in manganese accumulation that was higher in HeLa cells as compared to HAP1 cells. This is reflected by the ^55^Mn/^31^P ratio, which showed manganese enrichment levels in cells comparable to cerebral cortex and liver of Slc30a10^-/-^ mice (Fig. 1C, top panel) (Mercadante et al., 2019). However, both treatment doses lead to a 50-fold increase in manganese content in each cell type, comparable to liver of Slc30a10^-/-^ mice (Fig. 1C bottom panel), without changing the content of ^56^Fe, ^66^Zn or ^44^Ca in cultured cells or brain (Fig. S1C). These results indicate that the *in vitro* treatment conditions reproduce manganese levels attained in tissues of a mouse model of hypermanganesemia.

We reasoned that the dysregulated persistence of RNA transcripts of complementary sequence belonging to the H and L-strands could lead to dsRNA accumulation in mitochondria. We tested this hypothesis by detecting dsRNA co-localization with the mitochondrial RNA granule component GRSF1 by immunofluorescence with the J2 antibody, a monoclonal antibody that binds to dsRNA molecules larger than 40bp irrespective of sequence (Schonborn et al., 1991) (Fig. 1D). Both cell lines showed increased levels of dsRNA in mitochondria following 24h manganese treatment. dsRNA immunoreactivity was downregulated by inhibition of the mitochondrial RNA polymerase POLRMT with IMT1 (Bonekamp et al., 2020), indicating that the mitochondrial origin of the dsRNA signal induced by manganese is mitochondrial transcription (Fig. 1E). RNAse III was used as a control to selectively degrade dsRNA prior immune labeling (Fig. 1E).

### Manganese induced dsRNA activate type I Interferon responses

dsRNA of mitochondrial origin are known to escape from mitochondria into the cytoplasm where they are recognized by the dsRNA receptors, retinoic acid inducible gene 1 (RIG-1) and melanoma differentiation-associated protein 5 (MDA5) (Dhir et al., 2018; Chen and Hur, 2022). Once bound to dsRNA, these receptors are recruited to a multiprotein aggregate comprised of mitochondrial anti-viral signaling protein (MAVS) and signaling proteins. This multiprotein assembly activates the kinase TBK1, which phosphorylates IRF3, enabling its nuclear translocation to induce IFNβ transcription, thereby activating a cascade of interferon-stimulated genes (ISGs) (Fig. 2A) (Chan and Jin, 2022; Chen and Hur, 2022). To test the activation of this dsRNA receptor pathway by manganese, we expressed a reporter construct in which the IFNβ promoter drives luciferase expression. While HAP1 cells accumulated significant levels of dsRNA in mitochondria (Fig. 1D, E), no manganese concentration induced luciferase expression in these cells (Fig. 2B). This outcome suggests that dsRNA is retained in mitochondria in HAP1 cells, because the pathway was readily triggered by delivery of the membrane-permeable synthetic dsRNA agonist, poly(I:C) to the cytosol (Fig. 2B). In contrast, sub-IC_50_ manganese doses were sufficient to increase luciferase activity up to four-fold in HeLa cells (Fig. 2B). We next identified the receptor that recognizes manganese-induced mitochondrial dsRNA in the cytosol by knockdown experiments. Reduced expression of MDA5, but not RIG1, downregulated manganese-induced luciferase activity by 40% (± 11% SEM) (Fig. 2C). Combination knockdown of RIG1 and MDA5 did not change the inhibitory effect of MDA5 downregulation, indicating that of the two dsRNA receptors, only MDA5 is required to recognize manganese-induced dsRNA. In contrast, RIG1 was the receptor for poly(I:C) (Fig. 2C). We further confirmed mitochondrial dsRNA signaling by pharmacological inhibition of predicted components in the pathway (Fig. 2A, D). IFNβ promoter transcription was reduced by downregulating mitochondrial transcription with the inhibitor IMT1, which targets the mitochondrial RNA polymerase (POLRMT). Interferon promoter transcription was also reduced by interfering with dsRNA export to the cytosol by either inhibiting the permeability transition pore function in the inner mitochondrial membrane with cyclosporine A (CspA) or by blocking the Bax-Bak mitochondrial pore in the outer mitochondrial membrane with V5 peptide (Fig. 2A) (Dhir et al., 2018; Arnaiz et al., 2021; Luna-Sanchez et al., 2021). None of these inhibitors reduced manganese uptake by cells (Fig. 2D top panel) but all reduced IFNβ transcription by at least 25%, with 100% decrease when TBK1 was inhibited with GSK8612 (Fig. 2D bottom panel). Another potent activator of the antiviral pathway is the release of mitochondrial DNA (mtDNA) into the cytosol which activates cGAS-STING converging to TBK-1 activation (Fig. 2A). We excluded the contribution of mtDNA to signaling by pharmacological inhibition of STING with H151, which did not affect manganese or poly(I:C) induced luciferase transcription (Fig. 2E). In contrast, H151 effectively blocked STING activation-induced transcription by the agonist 2’3 c-di-AM(PS)_2_ Rp,Rp (ADU S100) (Fig. 2E). Taken together, these results indicate that manganese increases dsRNA in mitochondria of both cell lines, but dsRNA reach the cytoplasm where they bind to MDA5 and lead to increased IFNβ transcription only in HeLa cells.

**Figure 2:**
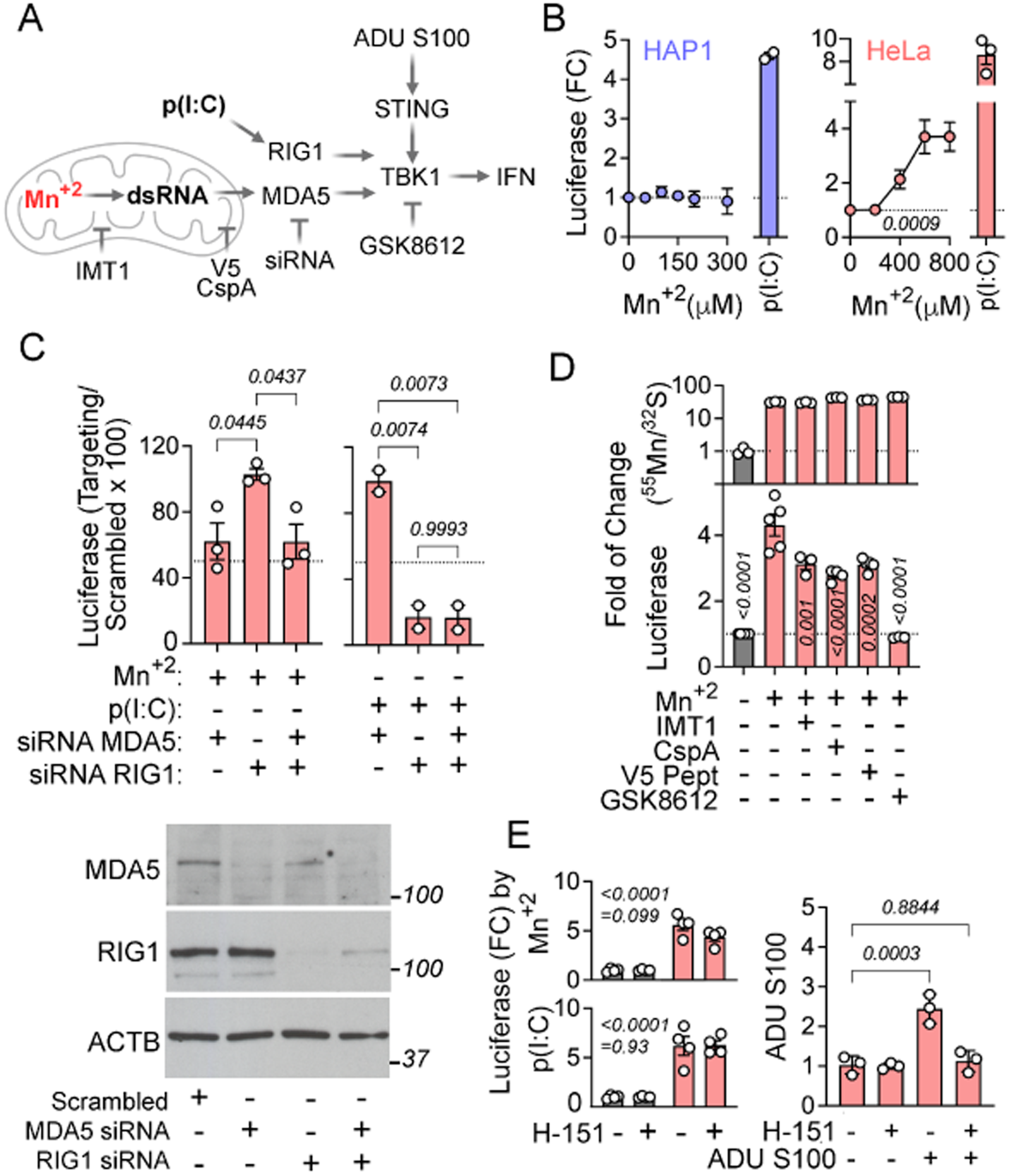
Manganese-induced dsRNA upregulates interferon β transcription. a) Diagram created with elements of Biorender, summarizing the pathway activated by manganese effects in mitochondria leading to IFNβ transcription. Depicted are the components targeted by inhibitors and the pathway agonists, poly (I:C) and ADU S100. b) IFNβ promoter luciferase reporter activity measured in either HAP1 or HeLa cells following 24h treatment with the indicated manganese dose. For positive control 10µg/ml of high molecular weight poly (I:C) was added. Average ± SEM of 3 experiments. One-way ANOVA (HAP1: F (5, 6) = 0.2137; HeLa: F (4, 10) = 11.62) followed by Dunnett’s multiple comparisons test. Dotted line marks luciferase activity in the absence of manganese. c) IFNβ promoter luciferase reporter activity measured in HeLa cells transfected with non-targeting and targeting siRNAs followed by 24h treatment with 800µM manganese or 10 µg/ml poly (I:C). Data are depicted as percentage of signal in scrambled sequence siRNA transfected cells. Average ± SEM of 3 experiments. Dotted line marks 50% reduction. One-way ANOVA (Manganese: F (2, 6) = 6.699; poly I:C: F (2, 3) = 46.64) followed by Tukey’s multiple comparisons test. Bottom panel depicts western blots of protein level downregulation following siRNA transfections. d) Top panel depicts manganese content determined by ICP-MS in HeLa cells incubated with 800µM manganese with and without the indicated inhibitor, identical conditions as employed for the bottom panel experiments. Bottom panel shows luciferase activity measured in HeLa cells following 24h treatment with the indicated manganese dose. For positive control, 10µg/ml of poly (I:C) were added. Average ± SEM of 5 experiments. One-way ANOVA (F (5, 20) = 63.05) followed by Bonferroni’s multiple comparisons test. e) Luciferase activity measured in HeLa cells following 24h treatment with 800µM manganese (top left panel) or 10µg/ml poly(I:C) (bottom left panel) in the presence or absence of the Sting inhibitor H-151. Right panel shows 0.5µM H-151 inhibition of luciferase induction by transfection of 1µM ADU S100 Sting agonist. Average± SE. Two-way ANOVA (Manganese: F (1, 12) = 133.2; drug: F (1, 12) = 3.191) followed by Bonferroni’s multiple comparisons test. Top numbers show manganese effect, bottom number show drug effect.

### Manganese-induced accumulation of mitochondrial dsRNA results in inflammatory responses

Next, we asked whether IFNβ transcriptional activation occurs in the context of a broader inflammatory response by profiling conditioned media of HAP1 and HeLa cells following a 48h manganese exposure with a cytokine profiling antibody array (Fig. 3A). While no significant changes were detected in HAP1 cells (Fig. S2B), we identified several cytokines and chemokines induced in response to increased manganese concentration in HeLa cells (Figs. 3A, B; Fig. S2A). Notably, the chemokines CXCL10 and GM-CSF were robustly upregulated by manganese, both of which have been shown to be upregulated following poly(I:C) treatment of epithelial cells (Choi et al., 2022). We detected modest increases in IL8, but not IL6, in the cytokine array which we confirmed by ELISA assay (Fig. 3C, D). We focused on IL8 production to monitor manganese-induced inflammatory responses as this cytokine is responsive to both interferon and NFκB activation by indirect and direct mechanisms, respectively and all cell types in the brain express receptors for this chemokine (Shkundin and Halaris, 2024).

**Figure 3:**
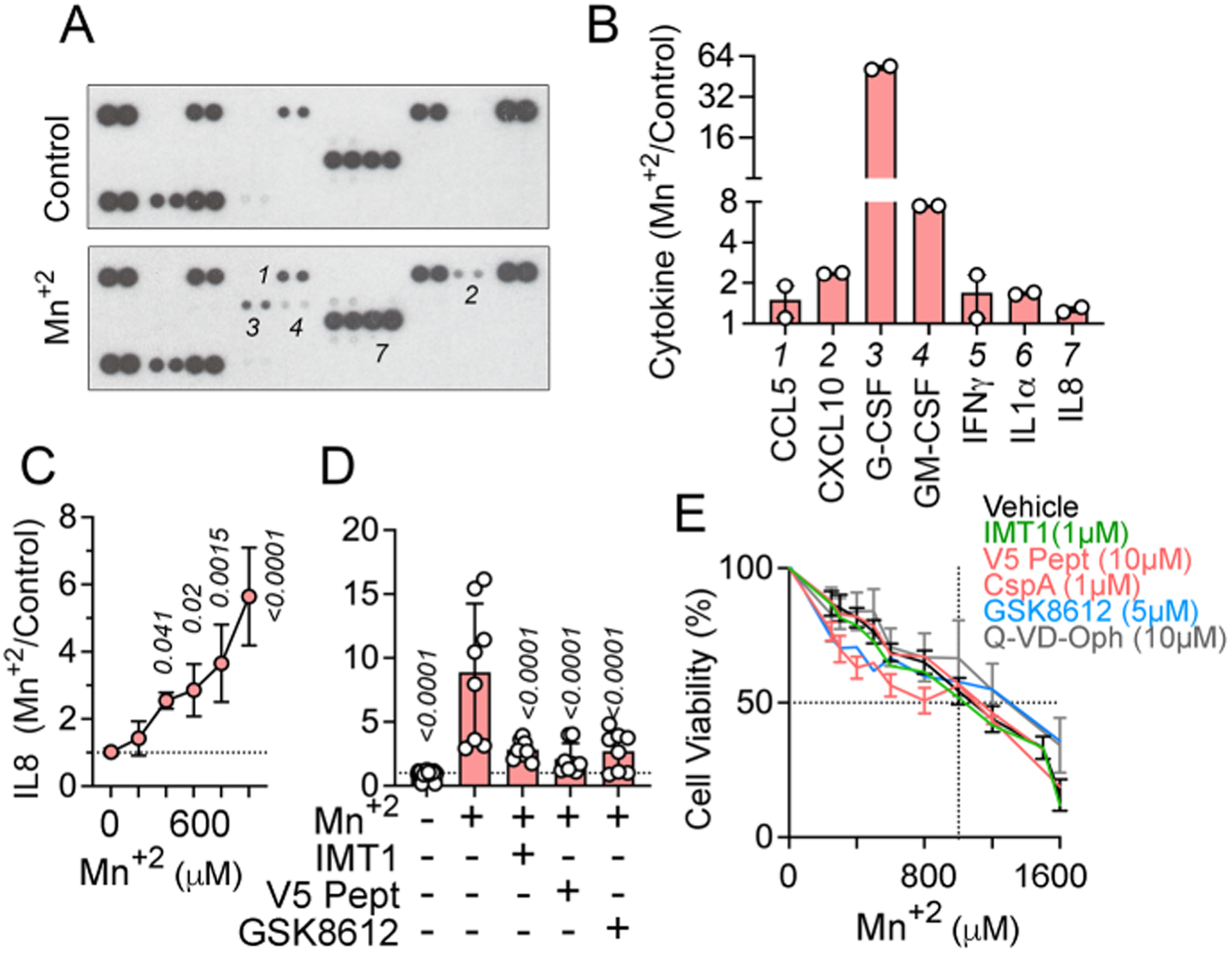
Manganese upregulates inflammatory cytokine secretion. a) Inflammatory cytokine antibody array probed with media conditioned for 48h collected from treated HeLa cells without or with 750µM manganese. b) Densitometry quantification of positive spots in the array blots shown in (a) and Fig S2A. c) IL-8 cytokine levels measured by ELISA in 48h conditioned media collected from HeLa cells treated without or with the indicated manganese concentration. Average ± SEM of 3 experiments. One-way ANOVA (F (5, 18) = 15.26) followed by Dunnett’s multiple comparisons test. d) IL-8 cytokine levels measured by ELISA in 48h conditioned media collected from HeLa cells treated without or with 800µM manganese and the indicated inhibitor. Average ± SEM of 4 experiments. One-way ANOVA (F (4, 35) = 72.35) followed by Dunnett’s multiple comparisons test. e) Cell survival of HeLa cells exposed to increased concentrations of manganese with and without the indicated inhibitor. Average ± SEM of 3-4 experiments. Two-way ANOVA (Manganese: F (10, 220) = 83.85; drug: F (5, 220) = 3.212) followed by Bonferroni’s multiple comparisons test.

Manganese induced IL8 expression following 48h treatment with a sub-IC_50_ manganese dose (Fig. 3C, E). This upregulation was reduced by inhibition of the mitochondrial RNA polymerase POLRMT with IMT1, the Bax-Bak pore function with V5 peptide, or a TBK1 inhibitor (Fig 3D). Importantly, none of the inhibitors affected cell survival of either cell line in response to manganese, which suggests that the inflammatory responses induced by dsRNA do not play a role in cytotoxicity (Fig. 3E, Fig. S2C). Addition of the pan-caspase inhibitor Q-VD-Oph reduced cytotoxicity in HAP1 cells (Fig. S2C) but did not affect HeLa survival, indicating that apoptotic mechanisms do not play a role in this inflammatory response either (Fig. 3E, Fig. S2C).

To better understand how manganese cytotoxicity and inflammation take place in a model system for brain tissue, we studied the effects of manganese on the mitochondrial transcriptome and cytokine secretion in human iPSC-derived cerebral organoids. We chose the unguided method for organoid growth as it provides diversified cell types and developmental paths to interrogate (Lancaster et al., 2017). To select the appropriate developmental timepoint to analyze, we tested for processing of mitochondrial transcripts during organoid development time with the MitoString panel, which we customized by adding probes to monitor differentiation of neuroprogenitor cells (NPC), neurons, and astrocytes. We found that neuroprogenitor cell markers declined overtime from day 30 on, with a steady increase in neuron maturation and astrocyte markers at days 60 and 100 (Fig. 4A, B). These transcriptional differentiation responses occurred concomitantly with morphological maturation and layering of the organoids as determined by confocal immunomicroscopy with antibodies against the cortical layer markers CTIP2 and SATB2 and by the progressive loss of immunoreactivity of the neuroprogenitor marker SOX2 (Fig. S3A).

**Figure 4:**
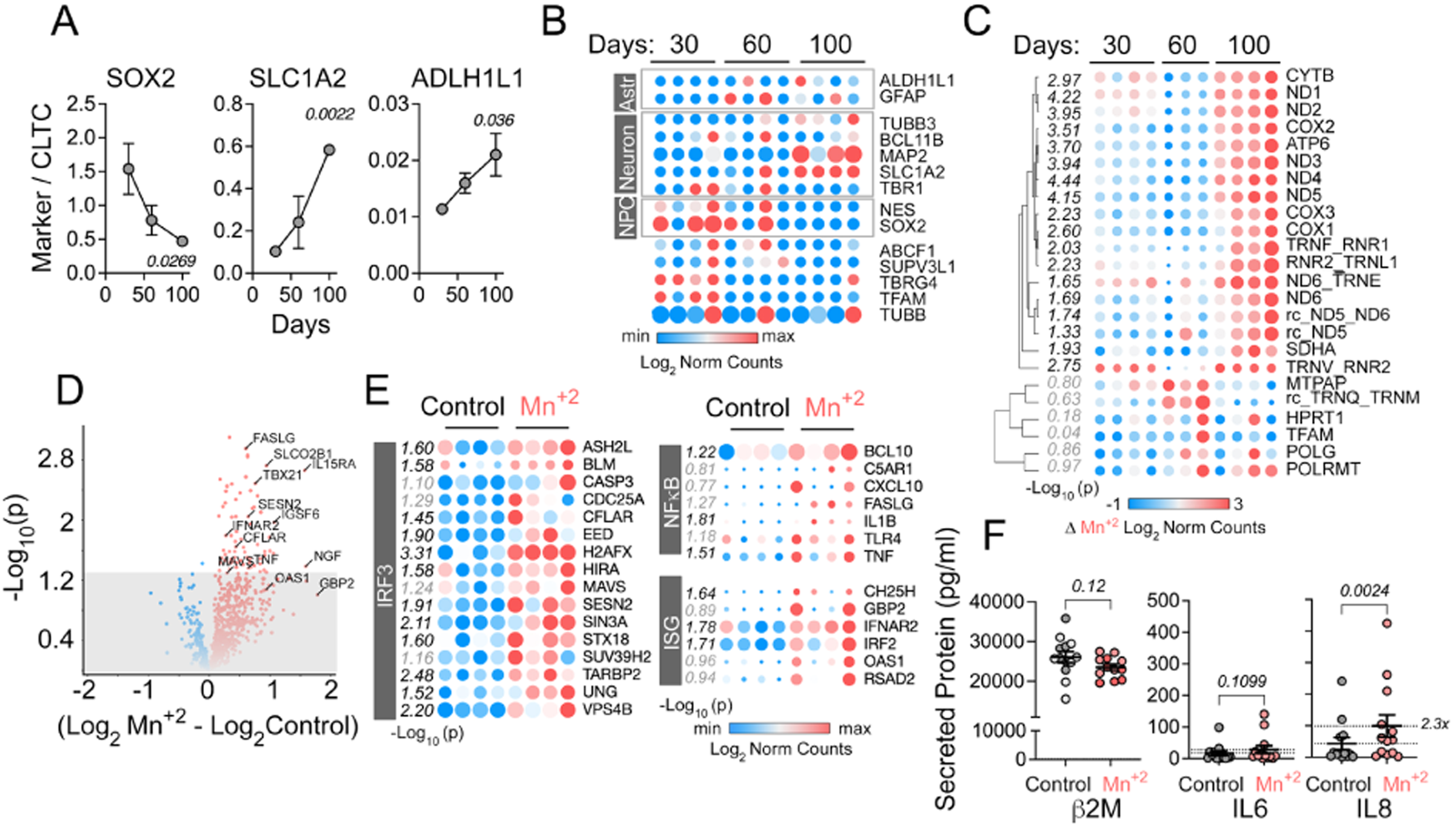
Manganese alters mitochondrial transcripts and induces the interferon type I response in human cerebral organoids. a) Lineage marker expression measured by NanoString over developmental time (Supplementary Dataset 1). Average ± SEM of 4 organoids per time point. One-way ANOVA (SOX2: F (2, 9) = 4.768; SLC1A2: F (2, 9) = 11.47; ALDH1L1: F (2, 9) = 3.949) followed by Dunnett’s multiple comparisons test. b) Heatmap of RNA levels of organoid differentiation markers and nuclear encoded mitochondrial transcripts measured by NanoString in individual organoids over time of development (Supplementary Dataset 1). Log_2_ of normalized counts of the gene were used for the heatmap. c) Heatmap of nuclear and mitochondrial encoded mitochondrial transcripts measured by NanoString in individual organoids over time of development (Supplementary Dataset 1). The difference of normalized counts from manganese-induced minus non-treated organoids per time point is expressed as Log_2_. Non-responsive organelles at 30 days were compared to responses in 100-day organelles. Two-tailed t-test expressed as −Log_10_. d) Volcano plot of the genes in the Neuroinflammatory panel that change expression in the transcriptome of 100-day organoids following 48h of 250μM manganese treatment (Supplementary Dataset 2). Significance threshold was set at 0.05 (marked by the line). e) Heatmap of IRF3 or NFκB transcriptional target genes and interferon stimulated genes included in the Neuroinflammatory panel (d) that change following manganese exposure (Supplementary Dataset 2). Two-tailed t-test expressed as −Log_10_. f) Cytokines in conditioned media collected from 100-day organoids following mock or 48h treatment with 250µM manganese. IL6, IL8 and beta 2 microglobulin levels were determined by MesoScale assay. Average ± SEM. Each dot represents an individual organoid. Data from two organoid batches. Statistics for beta 2 microglobulin paired t test and for IL6 and IL8 Wilcoxon matched-pairs signed rank test. For b, c and e symbol size is proportional to the Log_2_ value in between rows.

The organoids were treated with manganese for 48h at several times during development. After treatment, we collected organoids for RNA analysis and conditioned media for cytokine measurement. We used a dose of 250µM manganese, a concentration that is approximately equivalent to the metal concentrations estimated to accumulate in the striatum and globus pallidus of manganese-exposed primate and rodent models (Erikson et al., 2004). Using this strategy, we found that 30- and 60 day organoids did not show significant changes in mitochondrial-encoded RNAs or levels of their processing intermediaries in response to manganese. However, we found robust changes in 100- day organoids, including increases in most of the mitochondrial-encoded mRNA, some tRNA-mRNA junctions and several non-coding RNA encoded in the L strand of the mitochondrial genome (Fig. 4C). These modifications of the mitochondrial transcriptome occurred without changes in the nuclear-encoded mitochondrial genes TFAM, POLG, and POLRMT, excluding increases in mitochondrial content in 100-day organoids after 48h of manganese exposure.

We comprehensively interrogated the bulk 100-day organoid transcriptome to characterize the inflammatory responses induced by manganese employing the Neuroinflammatory (Fig. 4D, E) and Neuropathology (Fig. S3B, C) panels from NanoString. Hybridization with the Neuroinflammatory panel demonstrated upregulation of multiple inflammation-annotated mRNAs as represented in a volcano plot (Fig. 4D). Several genes that are transcriptional targets of IRF3 and NFκB as well as interferon stimulated genes (ISG) were significantly upregulated after manganese exposure (Fig. 4E). We measured secretion of inflammatory cytokines by analyzing the presence of IL6 and IL8 in conditioned organoid media with Mesoscale ELISA assays. Manganese significantly increased IL8 levels in conditioned media of organoids exposed to 250μM dose in two independent organoid cohorts (Fig. 4F, Fig. S3D). We observed a fraction of organoids with higher levels of basal cytokine secretion in the absence of manganese treatment (Fig. 4F, Fig. S3D). Basal untreated and manganese-induced cytokine secretion levels remained unaltered after normalizing organelle secretory activity and size by beta-2 microglobulin, a protein constitutively expressed and secreted by all cells (Fig. 4F). Analysis of the Neuropathology panel demonstrated that at this time of development and manganese concentration, there were no changes in the expression of transcripts annotated to either neurons and astrocytes or associated with disease mechanisms such as hypoxia, the unfolded protein response, or NRF2-dependent pathway signaling (Fig. S3C). Taken together, these results indicate that the manganese-induced mitochondrial and interferon type I inflammatory responses are present in brain cell types and are not part of an overt stress or cytotoxic response in brain organoids which lack microglia.

### Manganese induces inflammatory responses in astrocytes

To identify the cell types engaged in inflammatory responses to manganese in the organoids, we performed 10X single cell RNA sequencing (scRNAseq) on 12 individual 100-day organoids. Six organoids were left untreated and six were exposed to 250μM manganese for 48h. A total of 119,996 cells were analyzed and clustered into 8 distinct cell populations, including proliferative, fibroblast, choroid plexus, radial glia, astrocyte, neural progenitor, and two neuronal categories (Fig. 5A, QC metrics in Fig. S4A). All cell populations were proportionally present in all 12 organoids regardless of organoid size and manganese treatment (Fig. 5B, Fig. S4C). Cell clusters were manually identified based on differential gene expression profiles (Fig. 5C) and were of restricted expression as demonstrated by featureplots (Fig. S4B). Manganese did not preferentially affect the cell count of a specific cluster (Fig. S4D). We also examined the expression of selected inflammatory genes and identified astrocytes as the major cell population driving inflammatory responses in the organoids following manganese treatment (Fig. 5C). This response did not correlate with a specific enrichment in genes with functions in mitochondrial transcription or processing (Fig. S4E) genes involved in cellular manganese uptake or efflux, or metallothioneins (Fig. S4F). To gain insights into the mechanisms and molecular phenotypes elicited by manganese treatment in astrocytes, we obtained the differentially expressed genes via pseudobulking and DESeq2 to compare control and treated organoids (Fig. 5D), then conducted gene ontology analysis (Fig. 5E). Significantly up- and down regulated genes are represented in a volcano plot, where we highlight genes relevant to astrocyte function, CXCL10 and selected genes annotated to the pathways identified in bioinformatic analysis (Fig. 5D). Transcripts upregulated by manganese were enriched in genes annotated to interferon alpha responses, cholesterol biosynthesis and oxidative phosphorylation pathways in the MSigDB Hallmark 2020 database (Liberzon et al., 2015). Downregulated transcripts were annotated to UV response, hypoxia and TGFβ signaling (Fig. 5E). These manganese-induced responses were elicited across the treated individual organoids (Fig. 5F). Manganese increased the expression of CXCL8 and CXCL10, mitochondrial genes, subunits of the respiratory chain, and interferon response genes. Genes induced by manganese correlated with higher mRNA levels of astrocytic reactivity markers, cholesterol synthesis, and neuron supporting genes, but lower levels of transcripts annotated to phagocytosis (Fig. 5F). Feature maps of CXCL8 and CXCL10 expression revealed that a subpopulation of astrocytes expressed these chemokines, suggesting heterogeneity among the astrocyte responses (Fig. 5G). To characterize and identify unique attributes of these chemokine-expressing astrocytes, we subclustered astrocytes into 3 sub-populations (Fig. 5H, Astro 1, 2 and 3), where Astro 2 concentrated the cytokine expressing cells (Fig. 5G). Comparison of the differentially expressed genes in response to manganese showed that Astro 2 cluster also concentrated cells with the most robust induction of the interferon response genes, accumulated mitochondrial-encoded transcripts, and displayed reduced expression of HSD17B10 and REXO2. HSD17B10 is a component of the mitochondrial RNA processing complex RNaseP (Holzmann et al., 2008) and REXO2 participates in mitochondrial dsRNA degradation (Fig. 5I and Supplementary Dataset S7) (Szewczyk et al., 2020). The heterogeneous astrocyte responses detected by scRNAseq was also evident by immunomicroscopy of manganese-treated organoids stained for dsRNA and GFAP (Fig. 5J-1). We found that 36% of GFAP-positive astrocytes were also positive for dsRNA after exposing organoids to manganese (Fig 5J-2). We confirmed that dsRNA signal colocalized with mitochondrial markers in iPSC-derived astrocytes exposed to the same manganese challenge as organoids (Fig. 5J-3). In contrast to astrocytes, the neuronal cell groups in the organoids upregulated genes involved in neurogenesis including PANTR1, FOXG1, NEUSOD6, MEF2C and NTS, and in stress responses including SCRG1, ATF3, ATF5 and MSRA and mitochondrial genes, including NDUFB11 and HSD17B10 (Fig. S5A), which was reflected in the categories of enriched genes resulting from gene ontology analysis of the combined lists (Fig. S5B). Many of the genes upregulated in the scRNAseq were also detected in our bulk RNA analyses with the NanoString panels (Fig. S5C).

**Figure 5:**
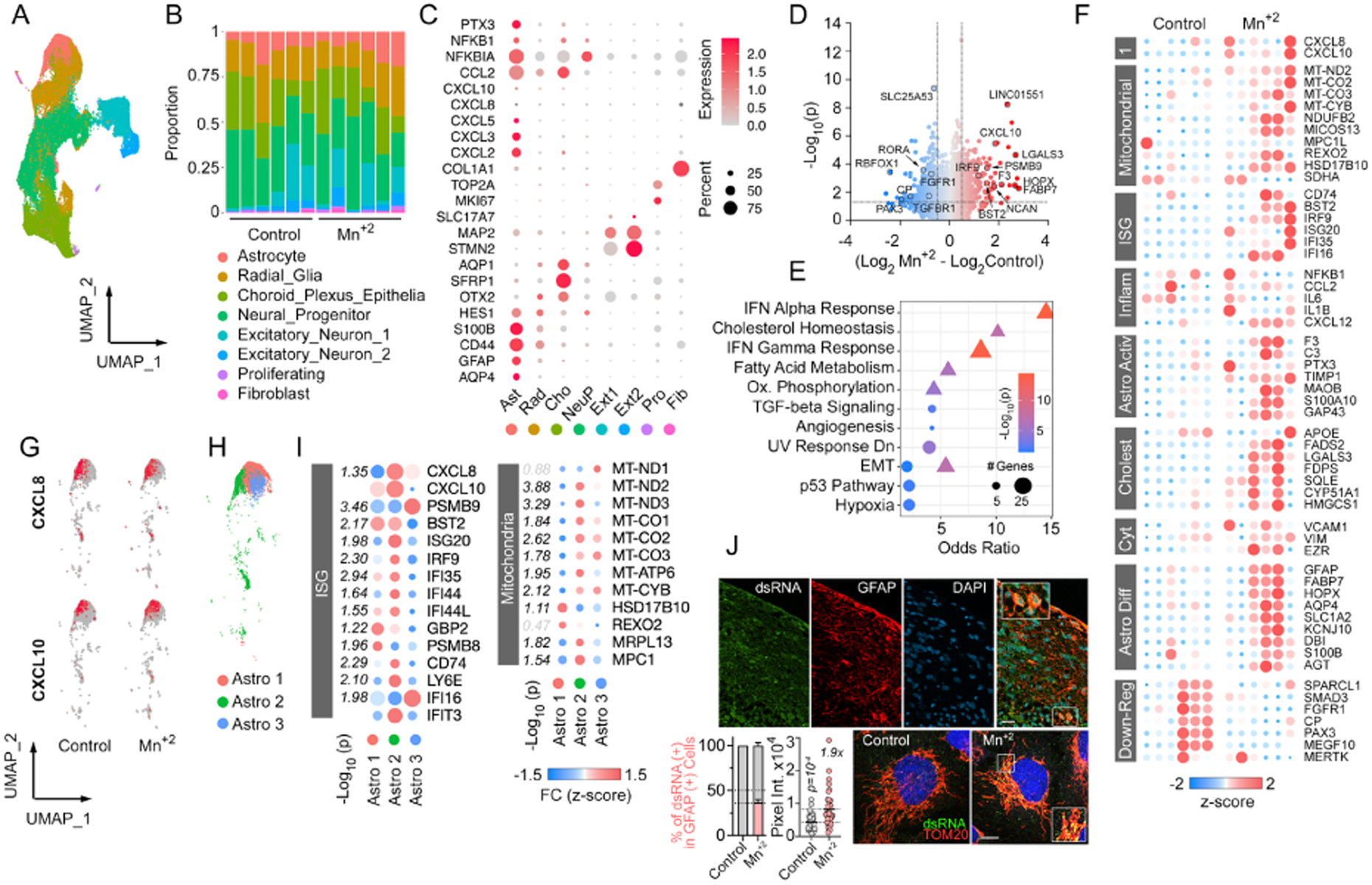
Single cell RNAseq analysis of manganese treated 100-day cerebral organoids. a) UMAP embedding of all samples (12 organoids aged 100 days) integrated and normalized with SCT transform. Cells are colored by cluster assignment. b) Proportion of cells assigned to each cluster from individual organoids. Color scheme is the same as in a). c) Dot plot of cell identity marker and inflammatory gene expression in cell clusters shown in a) (Supplementary Dataset 4). Dot size is proportional to percentage of cells expressing the gene and color reflects average expression level. d) Volcano plot of the manganese differentially regulated genes in astrocytes (Supplementary Dataset 5). Cut off is p<0.05 and 0.5< average log2FC <-0.5. Highlighted genes belong to the categories identified by gene enrichment analysis (e) or describing astrocyte function. e) Main up and down regulated pathways in manganese treated astrocytes when comparing the genes curated in Molecular Signatures Database employing EnrichR tool for gene ontology analysis. Triangles are up-regulated genes. Circles are downregulated genes. f) Heatmap of the scaled gene counts in the astrocyte cluster for individual organoids (Supplementary Dataset 6). g) Feature plot depicting the cells expressing CXCL8 and CXCL10 in control and manganese treated organoids. h) UMAP embedding of the subclustered astrocyte population from all organoids. i) Differential expression of genes belonging to the interferon response, mitochondrial transcripts and RNA processing in the three astrocytes subclusters in response to manganese (Supplementary Dataset 7). j) Immunofluorescence for dsRNA in manganese treated 100-day organoids and in iPSC derived astrocytes. 1) 100-day organoids stained for GFAP (red), DNA (blue) and dsRNA (green). Scale bar= 25μm. 2) Quantification of the percentage of GFAP cells that were positive for dsRNA in (1) and dsRNA in mitochondria in iPSC derived astrocytes shown in (3).1.9x refers to fold of increase, Mann Whitney test 3) Human astrocytes derived from iPSC treated or not with 250μM Manganese for 24h and stained for dsRNA (green) and the mitochondrial marker TOM20 (red). Scale bar =10μm.

### Chronic increased manganese induces inflammation in brain and liver in vivo

While organoids are an in vitro model for brain tissue well-suited to perform dose- and time-controlled experiments of acute manganese exposures, we asked next whether the reported phenotypes are present following chronic excess manganese accumulation in brain tissue in vivo. Mice with a global deficiency in the manganese efflux pump Slc30a10 progressively accumulate manganese in the brain from their diet (Mercadante et al., 2019). At eight weeks of postnatal age, Slc30a10^-/-^ mice reproduce several of the phenotypes observed in the human SLC30A10 genetic disease hypermanganesemia with dystonia 1 disorder (OMIM 613280), including motor deficits due to basal ganglia dysfunction (Mercadante et al., 2019). We asked whether cells in globus pallidus accumulate dsRNA in mitochondria, measured by immunofluorescence with J2 and colocalization with the mitochondrial marker Tom20. We detected increased dsRNA in GFAP-positive astrocytes (average of 35% +/- 20%SD in fields that had J2-positive cells) and in Nissl-positive neurons (average 50%+/-17% in fields that had J2-positive cells), partially co-localizing with the mitochondrial marker in globus pallidus of male and female Slc30a10^-/-^ mice (Fig. 6A and B). We were unable to detect dsRNA in microglia employing this approach. To test whether inflammatory responses are induced in this mouse model, we measured the expression levels of 32 cytokines in brain and liver samples from wild type and Slc30a10^-/-^ mice of both sexes as done previously (Wood et al., 2015). We included liver, which is known to accumulate the highest levels of manganese across tissues (Fig.1C and (Mercadante et al., 2019)). We found a complex pattern of altered cytokine expression modulated by sex, with higher expression levels of cytokines in KO tissues in both organs from male mice (Fig. 6C and Fig. S6A). To resolve cytokines whose expression was strongly associated with the mutant genotype, we analyzed these data by discriminant partial least squares regression analysis, which enables us to study multivariate datasets by reducing data from 32 measured variables to a reduced set of latent variables (LV1 and LV2) (Fig. 6D and Fig. S6B). Reducing the dataset to these latent variables allowed us to identify differences between genotypes distinguishing cytokine changes correlated with genotype from unrelated noise in the measurements (Geladi and Kowalski, 1986; Wood et al., 2015). Each LV represents a profile of cytokines that vary together between wild type and KO animals. We found that LV1 distinguished wild type animals and KO animals, irrespective of sex (Fig. 6D and Fig. S6B, x axis). We used a leave-one-out cross validation strategy, wherein each animal was iteratively left out, and the discriminant partial least squares regression analysis was re-computed. We found low standard deviation in each cytokine, suggesting that the involvement of each cytokine is not dependent on any single tissue sample (Fig. 6D and Fig. S6B). Moreover, the cytokine expression pattern was different in cortical and liver samples (Fig. 6E). In the brain, some of the upregulated cytokines are known to be produced by astrocytes (Giovannoni and Quintana, 2020), while Vegf, a cytokine responsive to hypoxia, was elevated in liver but not in cortex of Slc30a10^-/-^ animals, reproducing previous RNAseq findings (Prajapati et al., 2024b). In contrast, IL12b, was increased in both mutant tissues. (Figs. 6E and Fig.S6A). We profiled the wild type and Slc30a10^-/-^ brain cortex transcriptomes using a NanoString panel with 770 probes enriched for neuroinflammatory genes and glial reactivity. Seventy two genes were significantly upregulated when comparing wild type and knockout animals of both sexes combined (Fig. 6F), p<0.05 and fold change>1.5 (Dataset 9) with C3 and markers of the ISG among the most upregulated. The transcript alterations and cytokine profiles were more attenuated in female compared to male brains (Fig. 6G, H), however all mice exhibited upregulated interferon response and markers of reactive astrocytes and microglia (Fig. 6I).

**Fig 6:**
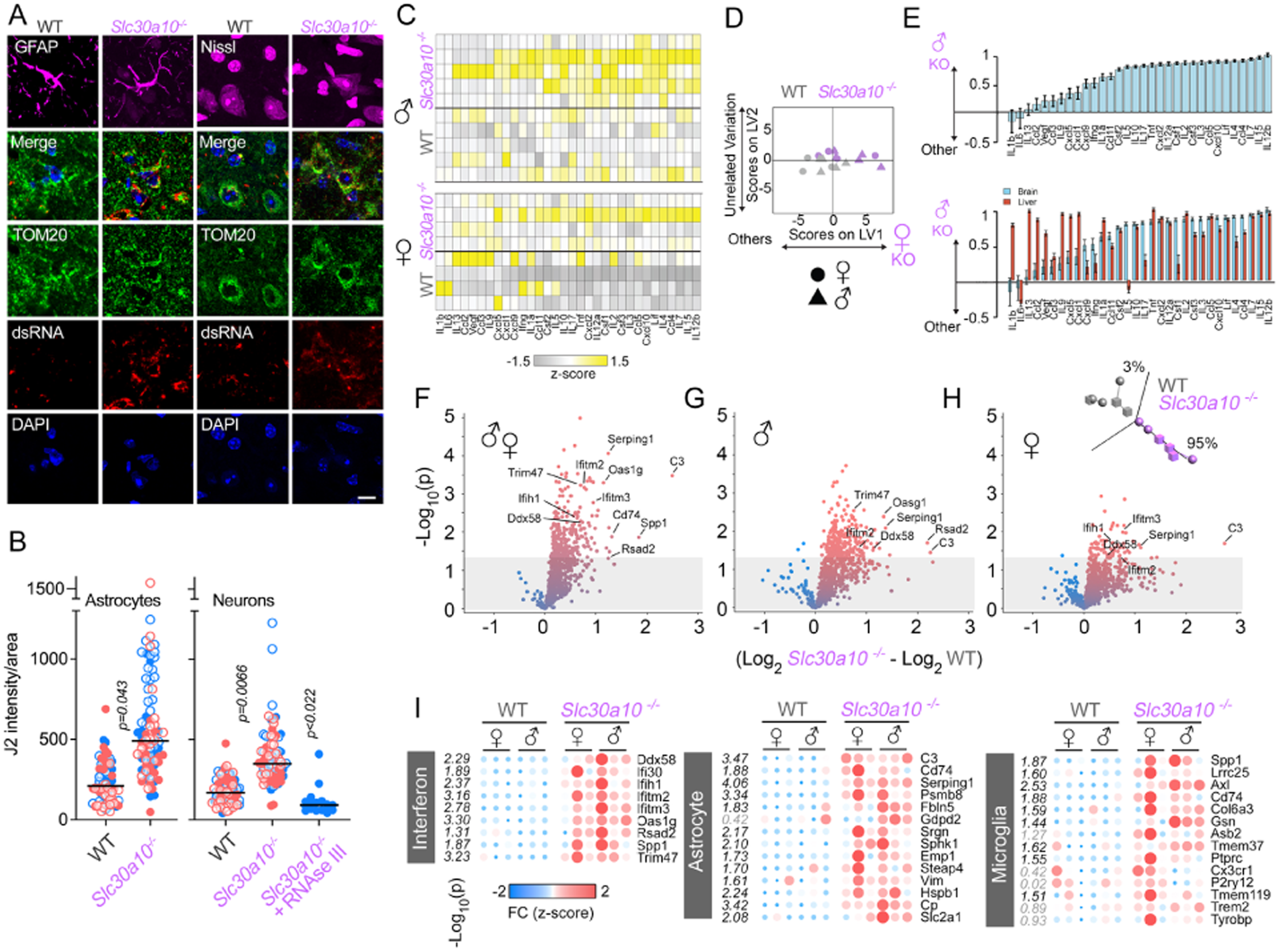
Brain from a mouse model of hypermanganesemia with dystonia 1 disorder has increased dsRNA accumulation and inflammatory responses. a) Immunofluorescence for dsRNA (J2), mitochondria (Tom20) and cell marker (GFAP or Nissl) in wild type and Slc30a10^-/-^ mouse brain. Scale bar= 10 μm. b) Quantification of J2 fluorescence intensity in 18-22 either GFAP- or Nissl-positive cells in the globus pallidus of 2 female (salmon circles) and 2 male (blue circles) wild type or Slc30a10^-/-^mouse. J2 signal reports dsRNA as it is downregulated by RNAse III treatment prior primary antibody incubation. The p value was calculated comparing considering number of cells and animals using SuperPlots. For the RNAseIII treatment the cells were compared to a wild type maile animal using a two-sided permutation t-test. c) Heat map of *z*-scored cytokine levels measured by Luminex multiplex assays in cortex samples of wild type and Slc30a10^-/-^ male and female mice. The cytokines are named by gene name. d) A discriminant partial least squares regression model constructed from the cytokine dataset regressed genotype. The model identifies a latent variable (LV1) that scores animals based on cytokine protein expression measurements and predicts genotype. LV2 describes variation that is not connected to genotype. LV1 and LV2 account for approximately 44% and 4% of the dataset variation, respectively. e) Top panel: LV1 is composed of cytokines that are elevated and able to predict the KO genotype in a leave-one-out cross validation in male mice samples (mean ± SD across LV1 generated for all models in the cross validation). Bottom panel: Comparison of LV1s from cortex and liver (in supplementary information) from Slc30a10^-/-^ male animals. f) Volcano plots of TBP normalized NanoString counts of RNA isolated from brain of a cohort of 6 wild type and 6 Slc30a10^-/-^ female and male mice, hybridized with probes targeted to neuroinflammatory gene sequences (Supplementary Dataset 9). g) Volcano plots of TBP normalized NanoString counts of RNA isolated from brain of a cohort of 3 wild type and 3 male Slc30a10^-/-^ mice. h) Volcano plots of TBP normalized NanoString counts of RNA isolated from brain of a cohort of 3 wild type and 3 Slc30a10^-/-^ female mice. Insert shows: linear near data reduction by principal component analysis (PCA) of all the RNA counts measured by the neuroinflammatory panel separates the samples of both sexes by genotype. i) Heatmaps of interferon stimulated genes, genes expressed in astrocytes or in microglia, included in the Neuroinflammatory panel that change significantly following manganese exposure. Two-tailed t-test expressed as −Log_10_.

Altogether, these results demonstrate the sustained expression of inflammatory responses with activation of type I interferon responses in Slc30a10^-/-^ brain tissue.

## Discussion

We identified a novel mechanism by which manganese induces inflammatory responses in non-immune cells, initiated by altered mitochondrial transcriptome processing that activates anti-viral type I interferon responses and inflammatory cytokines. Manganese causes dsRNA accumulation in mitochondria, detected by localization to the mitochondrial RNA granule in HeLa and HAP1 cell lines as well as in multiple brain cell types including astrocytes derived from human iPSCs, astrocytes in 100-day cerebral organoids, and in astrocytes and neurons in the globus pallidus of the Slc30a10^-/-^ mouse following manganese accumulation. Analysis of the mitochondrial transcriptome by NanoString demonstrated accumulation of unprocessed non-coding segments of mitochondrial transcripts in multiple cell types, including cell lines and cerebral organoids. Pharmacological and genetic interventions confirmed that mitochondrial dsRNA drives pro-inflammatory signaling by activating interferon responses in susceptible cell types such as HeLa cells and astrocytes. IFNβ transcription and IL8 production were reduced in HeLa cells by blocking either mitochondrial transcription with the mitochondrial RNA polymerase inhibitor IMT1 or dsRNA export to the cytosol with inhibitors targeting mitochondrial membrane permeability. We identified MDA5 as a cytoplasmic receptor sensing dsRNA and signaling to TBK1. Inhibitors or protein knockdowns targeting components of the pathway upstream of TBK1 activation, partially reduced responses but TBK1 inhibitors were highly effective, suggesting that manganese activates TBK1 through additional mechanism, possibly by binding to PKR or modulating ATM activity (Chan et al., 2000; Kim et al., 2018; Sui et al., 2022). Nonetheless, TBK1 inhibitors strongly downregulated IFNβ and IL8 without affecting cell survival, showing that this pathway drives inflammation disconnected from cell death.

We observed activation of type I interferon responses across complementary experimental models. Acute manganese exposure in cell lines and cerebral organoids produced phenotypes also observed in Slc30a10^-/-^ mouse brains, at an age when they show manganese accumulation and overt motor symptoms, indicating that these interferon responses may contribute to pathogenesis (Taylor et al., 2019). Phenotypic overlap across in vitro and in vivo models validated the manganese doses employed in vitro and the mechanisms identified. Because human brain manganese levels following exposures or in hypermanganesemia with dystonia 1 disease are not known, relevant exposure doses are inferred from animal model exposures, plasma levels after human exposure and findings in hepatic encephalopathy (Bowman and Aschner, 2014; Taylor et al., 2020). We argue that an important parameter to identify molecular mechanisms and signaling, is the intracellular manganese levels, which are defined by the balance of cell specific metal uptake and efflux, and organelle accumulation. We provide support for this premise by comparing the relative enrichment expressed as the manganese to phosphate ratio attained in acutely exposed cells, which matched the ratios in brain and liver of Slc30a10^-/-^ mice. Importantly, because manganese content of tissues represents the average of multiple cell types, it may be possible that some cell types in Slc30a10^-/-^ tissues may reach higher levels of manganese than the ones we report here.

Cerebral organoids modeled human astrocyte and neuron responses to manganese in vitro, in the absence of contributions to inflammation from microglia, immune cells and vasculature, indicating that astrocytes are sufficient to initiate inflammatory signaling. We show that changes in mitochondrial RNA transcript processing and inflammatory responses emerge in the organoids at the developmental time point of astrocyte maturation. Immunofluorescence confirmed that some of these astrocytes become GFAP-positive and accumulate dsRNA. Single cell RNAseq transcriptome analysis of organoids exposed to a short and non-lethal manganese treatment identified a small sub population of astrocytes as the main cell type driving IL-8 and CXCL10 expression/secretion and the induction of interferon response genes. In contrast, neurons induced stress responses reported by ATF3 and ATF5 expression without upregulating pro-inflammatory or interferon pathway genes. Staining of the globus pallidus of the Slc30a10^-/-^ mouse with J2 antibodies showed dsRNA accumulation in mitochondria in both cell types, suggesting that neurons may have a mechanism to prevent mitochondrial dsRNA pro-inflammatory activity. Evidence for one possible mechanism comes from one of the neuron clusters in organoids, which upregulated expression of suppressor of cytokine signaling protein SOCS1, which is part of a negative feedback mechanism to dampen inflammatory responses (Alexander and Hilton, 2004). Alternatively, dsRNA binding proteins in the neuronal cytosol, which are involved in cognition could be buffering dsRNA leaking from mitochondria, which in neurons, play important roles in cognition (Herbert, 2024). Neurons, particularly interneurons, express high levels of adenosine deaminase acting on RNA proteins (ADAR), which edit nucleotides to disrupt dsRNA structure (Lundin et al., 2020; Ansell et al., 2021; Dorrity et al., 2023; Shin et al., 2026)).

The transcriptional signature of the pro-inflammatory astrocytes in organoids shows a combination of deleterious markers such as reduced phagocytosis and high C3 expression, co-expressed with neuron-supporting phenotypes, such as cholesterol biosynthesis, cytoskeleton genes controlling cell surface area, and glutamate transporters. Multiple genes of this complex astrocyte signature were upregulated in Slc30a10^-/-^ mice cortical tissue indicating the presence of these mechanisms in the mouse model. In the cortex, microglial markers of reactivity were also upregulated; however, they did not fully overlap with a specific reactivity phenotype such as Disease Associated Microglia (DAM), defined by downregulation of homeostatic genes Cx3cr1, Tmem119 and P2y12 with upregulation of Trem2 and Tyrodp (Chen and Colonna, 2021; Paolicelli et al., 2022). Altogether, our findings suggest that astrocytes inducing the expression of C3 may be a primary manganese target, which together with microglial activation, is predicted to change synaptic function and pruning rates (Stephan et al., 2012) as well as promote neurodegeneration (reviewed in (Lawrence et al., 2023).

We show that type I interferon responses are triggered within 24-48h of manganese exposure. Our findings are supported by published transcriptome datasets, which reveal manganese-dependent induction of interferon responses in human fetal astrocytes treated for 7-days in vitro (Sengupta et al., 2007). Previous bulk RNA transcriptomic studies in Slc30a10^-/-^ mice showed upregulation of GFAP, CXCL10 and the interferon response genes Oas2 and 3 in brain, but not in liver (Prajapati et al., 2024a). Bulk RNAseq analysis of basal ganglia from 1 month old Slc30a10^-/-^ mice showed interferon responses as one of the major categories of upregulated genes in 2 of the 3 regions probed and could be reversed by lowering dietary manganese (Warden et al., 2024). Because astrocytes exhibit high manganese uptake and persistent activation into adulthood when mice are exposed as juveniles(Moreno et al., 2009), transient exposures may trigger lasting interferon responses. The contribution of this pathway to neuropathology could be compounded as all cells in the brain express IFNβ receptors and a basal tone is essential for development and cognition (Viengkhou and Hofer, 2023; Dorrity et al., 2024). Low astrocytic IFNβ promotes synaptic spine formation and cognitive function in hippocampus (Hosseini et al., 2020) and primes microglia phagocytic activity towards neuronal bodies early in development (Escoubas et al., 2024). Complete absence of this pathway or excess is pathogenic. For example, IFNβ or IFNR loss of function is sufficient to cause Parkinson’s- like neurodegeneration(Ejlerskov et al., 2015). Additionally, dystonia is associated with excessive interferon signaling in Aicardi Goutières syndrome (Peixoto de Barcelos et al., 2024) and is a secondary occurrence during Type I IFN therapy (Viengkhou and Hofer, 2023; Zhang et al., 2024). Thus, dysregulated IFNB release from manganese-exposed astrocytes may mediate cognitive and/or motor symptoms.

We previously identified mitochondrial transcripts processing as a new target for manganese, which we now show can become an effector of manganese-induced signaling to trigger inflammation. It remains unclear whether manganese is acting indirectly on cytosolic pathways that then affect mitochondrial RNA metabolism or directly on mitochondrial RNA or proteins. Manganese readily accumulates in mitochondria (Gavin et al., 1992), disrupting respiratory chain function by poorly understood mechanisms that lead to reduced oxygen consumption and increased ROS production in astrocytes (Sarkar et al., 2018), iPSC-derived dopaminergic neurons (Budinger et al., 2025) and in HAP1 cells ((Werner et al., 2022), reviewed by TE Gunter p591, DC Dorman 2023). Manganese could affect RNA processing or stability by direct interaction with phosphate groups of the RNA backbone affecting conformation and/or folding (Izatt et al., 1971), or by interfering with catalytic activity by displacing magnesium in enzymes that use this divalent cation as a cofactor ((Smith et al., 1992), (Brannvall and Kirsebom, 1999),(Foster et al., 2014)). Compared to nuclear mRNA, mtRNA is 160 times more abundant, but is less stable (McShane and Churchman, 2024). Given the elevated mtRNA output of a cell (30% of total RNA) compared to low translational output, dsRNA accumulation could occur without impacting other mtRNA functions such as translation and respiratory complex assembly. CRISPR screens demonstrate that functional interference of multiple mitochondrial enzymes can alter RNA processing and lead to dsRNA accumulation (Wolf and Mootha, 2014; Kim et al., 2024). Accumulation of some mitochondrial metabolites such as fumarate generated by the Kreb cycle enzyme fumarate hydratase (Hooftman et al., 2023) or itaconate generated by mitochondrial aconitate decarboxylase, (O’Carroll et al., 2024) promote dsRNA accumulation and inflammatory responses. Metabolic changes in the cytoplasm, such as hypoxia reduces dsRNA accumulation and interferon signaling by suppressing mitochondrial transcription in cancer cell lines (Arnaiz et al., 2021; Peng et al., 2021). Thus, manganese could be only one of many triggering mechanisms for mitochondrial dsRNA signaling.

## Acknowledgments

This work was supported by NIH grant R01ES034796 to AG and EW, Udall Center Pilot Project Grants for Parkinson’s Disease Research to AG, Hercules Pilot Award NIEHS P30ES019776 to EW and AG, NIH T32NS096050, HHMI Gilliam Fellowship, ARCS Foundation Scholar Award to HM-V, NIH K00 ES033033 and BWF PDEP to MMS, NIMH R01 MH125956 to SAS, 1R01AG075820 to LBW, FGRM was funded in part by the Cell and Tissue Engineering Training Grant NIH T32GM145735 NIGMS, NIH R01 DK110049 to TBB. Research reported in this publication was supported in part by Emory University subsidized cores: Emory Integrated Genomics Core (EIGC, RRID# iSCR_023529), Emory Multiplexed Immunoassay Core (EMIC), Emory Stem Cell and Organoids Core (ESCOC RRID# SCR_023264), Integrated Cellular Imaging (ICI, RRID:SCR_023534) of the Winship Cancer Institute of Emory University and NIH/NCI under award number 2P30CA138292-04.

## Data and code availability

All datasets are available through the Gene Expression Omnibus at GEO: GSE316010 All scripts generated will be available without restrictions upon request via the Sloan Lab GitHub.

