## Supplementary figures for "Mitochondrially Transcribed dsRNA Mediates Manganese-induced Neuroinflammation"

Hadassah Mendez-Vazquez *et al.*

**This PDF file includes:**

Materials
Figs. S1 to S4
Legends for Dataset S1 to S8
References

**Other Supplementary Materials for this manuscript include the following:**
Datasets S1 to S9

### Materials

| Reagent | Company | catalog number | RRID |
|---|---|---|---|
| iPSC | GIBCO | A18945 | CVCL_RM92 |
| HeLa | ATCC | CCL-2 | |
| HAP1 | Horizon | C631 | CVCL_Y019 |
| HAP1 SLC30A10KO | Horizon | HZGHC004693c005 | CVCL_TM87 |
| DMEM | Corning | 10-013-CV | |
| Iscove's DMEM | Corning | 10-016-CV | |
| OptiMEM | Invitrogen | 31985-070 | |
| Lipofectamine RNiMAX | Invitrogen | 13778-030 | |
| Embryonic Stem Cell qualified Matrigel | Corning | 354277 | |
| pNiFty3-L-Fluc-Puro | InvivoGen | pnf3p-fluc4 | |
| polyICHMWvLyoVec | InvivoGen | tlrl-piclv | |
| In Solution Q-VD-Oph, non O-methylated | Millipore Sigma | 551476 | |
| J2 | SCICONS | RNT-SCI 10010200 | AB_2922431 |
| GRSF1 | Abcam | EPR 16678 | AB_2827628 |
| GFAP | Chemicon | MAB360 | AB_11212597 |
| Tom20 (F-10) | SC | sc-17764 | AB_628381 |
| MDA5 | CST | D14E4 | AB_10694490 |
| Rig-1 | CST | D14G6 | AB_2269233 |
| Sox2 | R&D | AF2018 | AB_355110 |
| Tuj1 | Biolegend | 801201 | AB_2313773 |
| Ctip2 | Abcam | ab18465 | AB_2064130 |
| Satb2 | Abcam | ab34735 | AB_2301417 |
| RNAseIII | Ambion | AM2290 | |
| SUPERase-In Rnase inhibitor | Ambion | AM2696 | |
| ON-TARGETplus Non-targeting pool | Dharmacon | D-001810-10-05 | |
| ON-TARGET Human DDX58 siRNA SMART Pool | Dharmacon | L-012511-00-005 | |
| ON-TARGET Human IFIH1 siRNA SMART Pool | Dharmacon | L-013041-00-005 | |
| Chromium GEM-X Single Cell 3' Chip kit v4 | 10xGenomics | 1000690 | |
| Human Cytokine array | R&D Systems | ARY005B | |
| The human Proinflammatory 9-Plex panel | MesoScale | K15007B-1 | |
| R-PLEX Human B2M Antibody Set | MesoScale | F21AKZ-3 | |
| XT_Hs_Neuroinflamm_CSO | NanoString | 115000230 | |
| XT_Hs_Neuropath_CSO | NanoString | 115000200 | |
| XT_Mm_Neuroinflamm_CSO | NanoString | 115000237 | |
| V5 peptide inhibitor | Millipore Sigma | V7754 | |
| AZDye 546 Tyramide | VectorLab | CCT-1539-1 | |

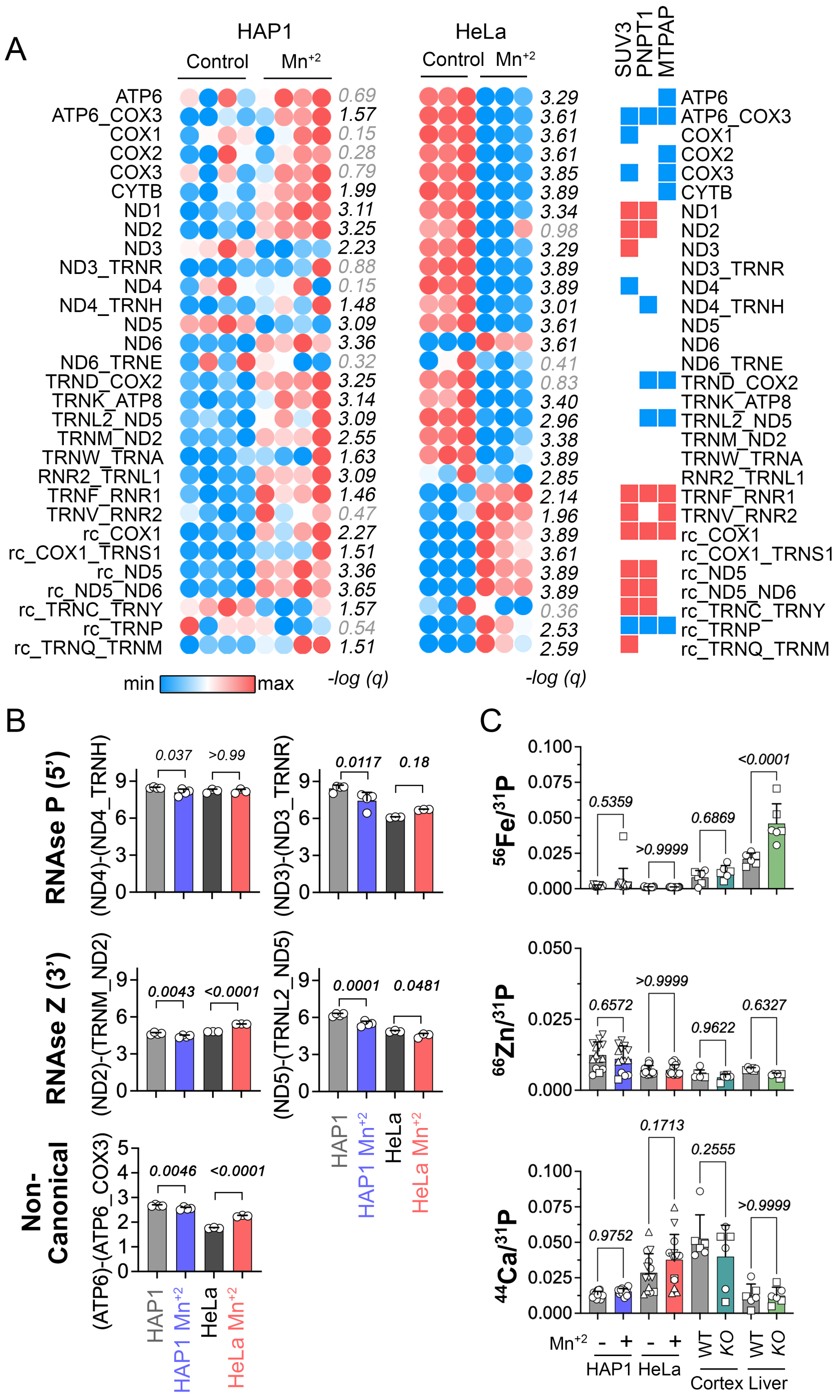


**Figure S1: MitoString panel of mitochondrial transcripts following manganese treatment and other relevant metal content of manganese treated cell lines and Slc30a10$^{-/-}$ mouse tissue**

a) Heat map of normalized $log_2$ counts of mRNA and junctions RNA detected by the MitoString panel in HAP1 and HeLa cells treated with manganese for 24h. Each replicate is shown with FDR corrected p value, significatively changed are highlighted in bold. For comparison, the right panel indicates the direction of change obtained when disrupting by CRISPR the mitochondrial enzymes

SUV3, PNPT1, MTPAP, adapted from (Wolf and Mootha, 2014), which suggests that all the changes induced by manganese can not be explained by deficient activity of one of these enzymes.

b) Comparison of non-canonical, 3' or 5' processing RNAse activity reported by the targeted mRNA/junction ratios of normalized $\log_2$ RNA counts in both cell lines without and with manganese treatment. Two-way ANOVA (ND4/ND4_TRNH Cell: F (1,10) = 0.6921; manganese: F (1,10) = 3.8; ND3/ND3_TRNR Cell: F (1,10) = 49.48, manganese: F (1,10) = 0.7634; ND2/TRNM_ND2 Cell: F (1,10) = 205.7, manganese: F (1,10) = 24.22; ND5/TRNL2_ND5 Cell: F (1,10) = 181.1, manganese: F (1,10) = 39.00; ATP6/ATP6_COX3 Cell: F (1,10) = 804.6, manganese: F (1,10) = 77.12 followed by Bonferroni multiple comparisons test.

c) Phosphate normalized $^{56}Fe$, $^{66}Zn$ and $^{44}Ca$ content of manganese treated cell lines and Slc30a10$^{-/-}$ mouse tissue. Metals were measured by ICP-MS at 24h post treatment in cell pellets of at least 3 independent experiments, each represented by different symbols, in brain cortex punch biopsies and liver of 8 weeks old wild type (WT) and Slc30a10$^{-/-}$ (KO) mice (3 female (○ ), 3 male ( □ ). No sex differences were noted following two-way ANOVA test.

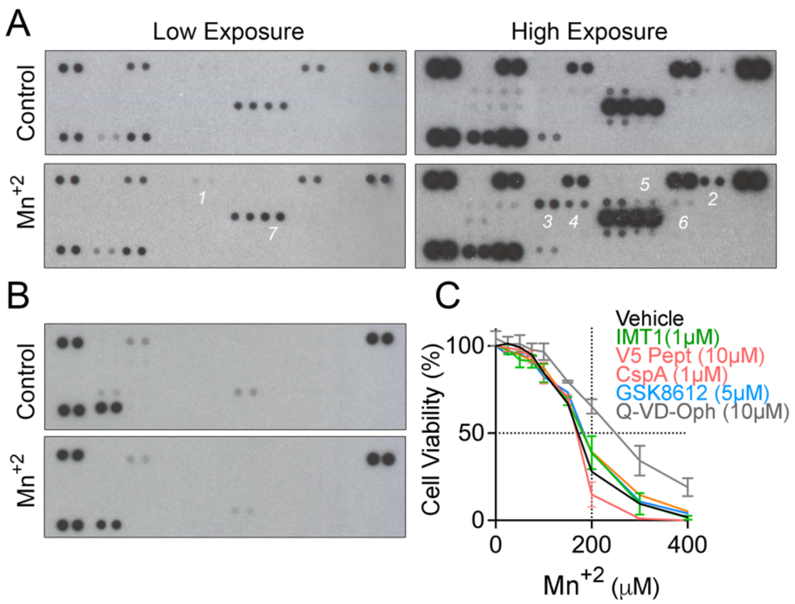


**Figure S2: Cytokine antibody array for HAP1 and HeLa, HAP1 survival to manganese in the presence of inhibitors**

a) Low and high exposures of the inflammatory cytokine antibody array probed with media conditioned for 48h collected from treated HeLa cells without or with 750µM manganese.

b) Inflammatory cytokine antibody array probed with media conditioned for 48h collected from treated HAP1 cells without or with 200µM manganese.

c) Cell survival of HAP1 cells exposed to increased concentrations of manganese with and without the indicated inhibitor. Average ± SEM of 3-4 experiments. Two-way ANOVA (manganese: $F(10, 236) = 93.62$; inhibitors: $F(6, 236) = 4.399$) followed by Bonferroni's multiple comparisons test.

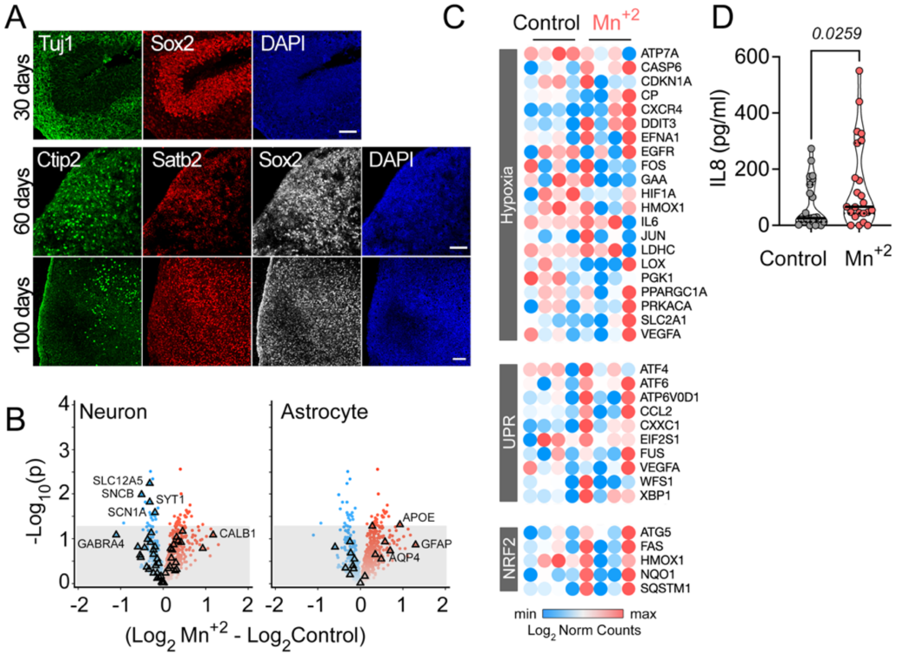


**Figure S3: Organoid characterization and Neuropathology NanoString panel analysis**

a) Morphological organoid characterization: Day 30 organoids were characterized for Sox2 positive neuroprogenitor cells organized into ventricular like zones surrounded by Tuj1 positive neurons. At 60 days, neuronal cells spatially segregated in regions as shown by immunohistochemistry for the expression of Satb2 (layer2-4) in cells close to the surface, with deeper located Ctip2 positive cells (layer 5 neurons) and Sox2 positive neuroprogenitors. At day 100, Ctip2 positive cells organize into cortical layers. Size bar= 50µm.

b) Volcano plot of the 100-day organoids following 48h of 250µM manganese treatment (Supplementary Dataset 3). Triangles mark all genes annotated as neuronal or astrocytic according to the PanglaoDB database[1] and present in the Neuropathology panel. Significance threshold was set at 0.05 (marked by the line).

c) Heatmap of $Log_2$ LARS-normalized counts of transcripts after mock or 48h treatment with 250µM manganese (Supplementary Dataset 3). Depicted are transcripts annotated to Hypoxia, the Unfolded Protein Stress responses, or NRF2 transcriptional target genes according to MSigDB Hallmark and included in the Neuropathology panel.

d) IL8 in conditioned media collected from an independent cohort of 100-day organoids following mock or 48h treatment with 250µM manganese. IL8 was measured by ELISA. Each dot represents an individual organoid n=23. Kolmogorov-Smirnov test.

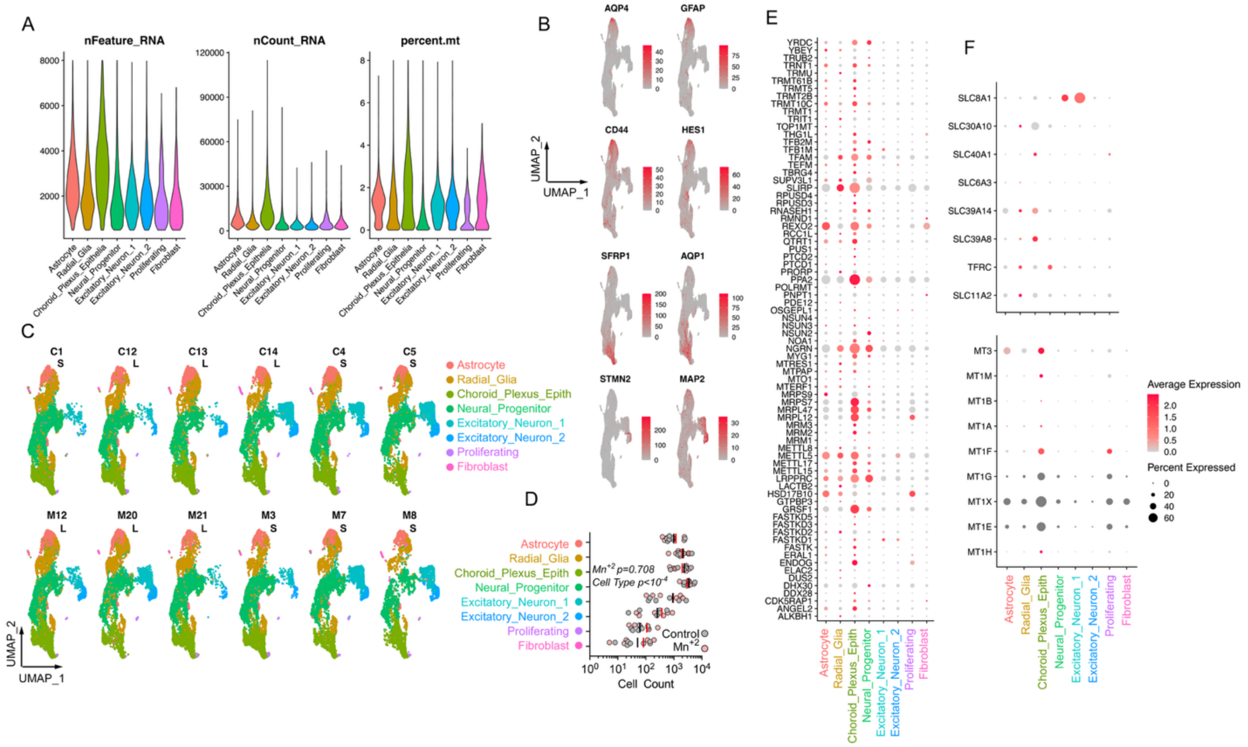

**Figure S4: Single cell QC, UMAPs broken down by metadata (organoid, treatment, size) and featureplots of major cell class markers**

a) Violinplots showing QC features: number of genes detected in each cell cluster (nFeature), number of RNA molecules detected in each cluster (nCount), and percentage of reads mapping to MT genes for each cell cluster.

b) Featureplots for major cell class markers: astrocytes (AQP4, GFAP, CD44, S100B), radial glia (HES1), choroid plexus (SFRP1, AQP1), neurons (STMN2, MAP2). Max cutoff at quartile 99.

c) UMAPs for each organoid non-treated (top row) or exposed to Mn (bottom row). Cells are colored for cluster assignment including astrocyte subclusters. S= small (2mm< diameter<2.5mm), L= large (2.6mm< diameter< 5mm).

d) Cell number counts for each cluster in each organoid in the absence and presence of manganese. Two-way ANOVA (cell type: $F(7, 80) = 0.1818$; manganese: $F(1, 80) = 0.1286$) followed by Šídák's multiple comparisons test with cell type and metal treatment as variables. Vertical black and red bars depict the mean cell count for control and manganese-treated organoids, respectively.

e) Dot plot of mitochondrial genes in the categories for mitochondrial transcription and RNA metabolism in MitoCarta 3 (Rath et al., 2021), expressed across cell clusters. Dot size is proportional to percentage of cells expressing the gene and color reflects average expression level.

f) Dot plots of genes for manganese efflux and uptake (Chen et al., 2018) (top panel) and metallothionein (bottom panel) expressed across cell clusters. Dot size is proportional to percentage of cells expressing the gene and color reflects average expression level.

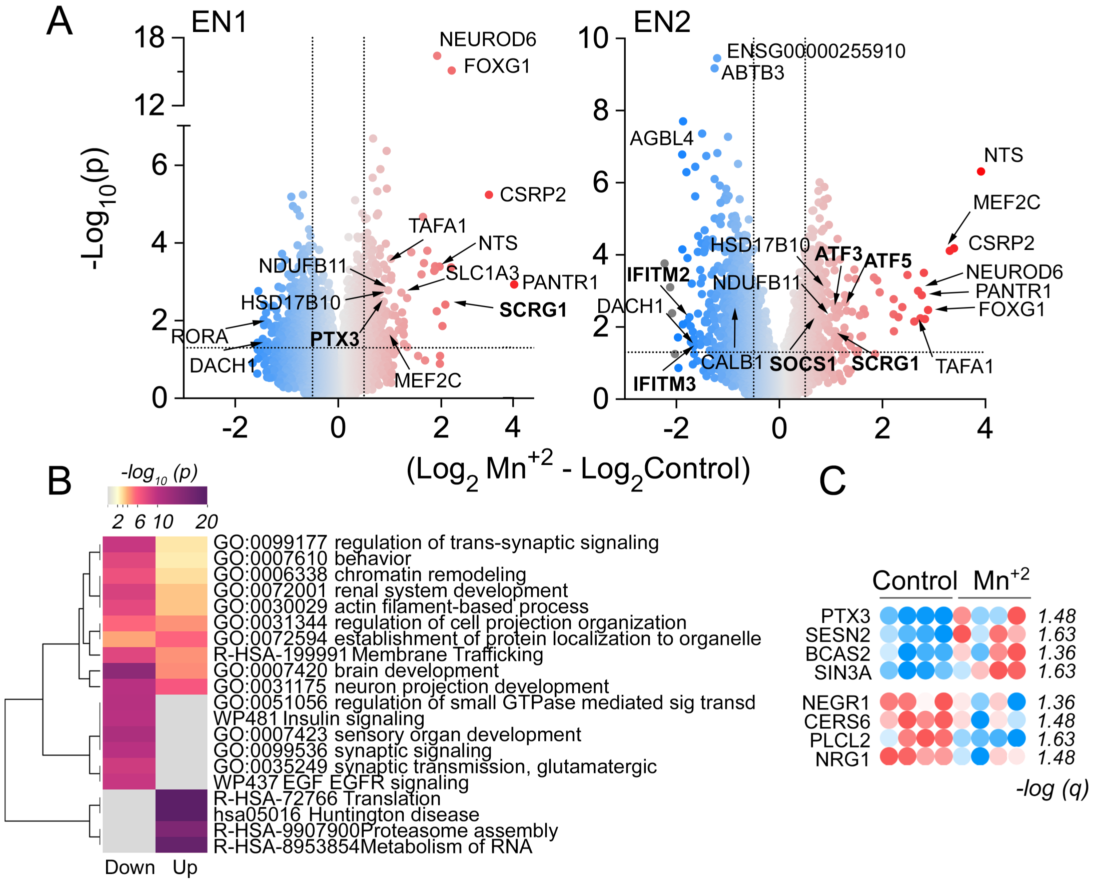


**Figure S5: Differential gene expression analysis of the neuronal clusters in manganese treated organoids**

a) Volcano plot of the manganese differentially regulated genes in two neuronal cell clusters (Supplementary Dataset 8). Cut off is $p<0.05$ and 0.5< average log2FC <-0.5. Highlighted genes are regulated by manganese in both cell clusters. In bold are highlighted genes regulated one of the populations only.

b) Main up and down regulated pathways in the neuronal clusters of manganese treated organoids when the differentially regulated genes were annotated and analyzed using the Metascape tool for gene ontology analysis (Supplementary Dataset 6) (Zhou et al., 2019). Triangles are up-regulated genes. Circles are downregulated genes.

c) Heatmap of $Log_2$ normalized counts of transcripts regulated by manganese in 100 Day organoids detected by Neuroinflammatory and Neuropathology NanoString panels (Supplementary Datasets 2 and 3) that overlap with genes in the neuronal subclusters measured in scRNAseq.

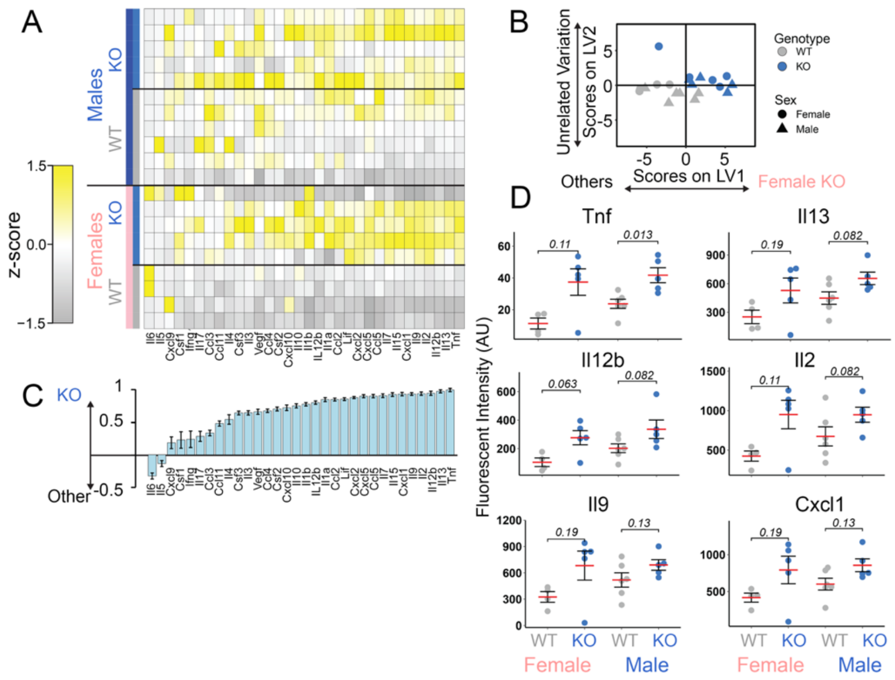

**Figure S6: Liver from a mouse model of hypermanganesemia with dystonia 1 disorder has increased pro-inflammatory cytokines**

a) Heat map of *z*-scored cytokine levels measured by Luminex multiplex assays in liver samples of wild type and Slc30a10$^{-/-}$ male and female mice. The cytokines are named by gene name.

b) A discriminant partial least squares regression model constructed from the cytokine dataset regressed genotype as in figure 6B. LV1 and LV2 account for approximately 43% and 9% of the dataset variation, respectively.

c) LV1 is composed of cytokines that are elevated and able to predict the KO genotype in a leave-one-out cross validation (mean ± SD across LV1 generated for all models in the cross validation).

d) Example cytokine levels in liver that yielded the highest LV1 scores.

### Legend for Datasets (separate file)

**Dataset S1** CLCT normalized counts from MitoString panel in organoids at 30, 60 100 days used in panels A and B in Figure 4.

**Dataset S2** TBP normalized counts from Neuroinflammatory panel in 100-day organoids used in panels E and D in Figure 4.

**Dataset S3** LARS normalized counts from Neuropathology panel in 100-day organoids used in panels B and C in Figure S3.

**Dataset S4** Genes used for cluster ID in Figure 5C.

**Dataset S5** ConVsMn.Astrocyte.DESeq.pseudobulk.allresults data used to generate the volcano plot in Figure 5D.

**Dataset S6** Gene expression levels used to generate heat map in Figure 5F.

**Dataset S7** Differentially expressed genes in astrocyte subclusters used to generate heat map in Figure 5I.

**Dataset S8** ConVsMn.DESeq.pseudobulk.allresults data for two neuron cell clusters used to generate the volcano plot in Figure S5A and ontology analysis in S5B.

**Dataset S9** TBP normalized $Log_2$ counts from Neuroinflammatory panel in brain cortex of wt and $Slc30a10^{-/-}$ mice used in panels D through G in Figure 6.
